# Regional cytoarchitecture of the adult and developing mouse enteric nervous system

**DOI:** 10.1101/2021.07.16.452735

**Authors:** Ryan Hamnett, Lori B. Dershowitz, Vandana Sampathkumar, Ziyue Wang, Vincent De Andrade, Narayanan Kasthuri, Shaul Druckmann, Julia A. Kaltschmidt

**Affiliations:** Department of Neurosurgery, Stanford University School of Medicine, Stanford, CA 94305 USA; Wu Tsai Neurosciences Institute, Stanford University, Stanford, CA, 94305, USA; Department of Neurobiology, University of Chicago, Chicago, IL 60637 USA; Biosciences Division, Argonne National Laboratory, Lemont, IL 60439 USA; Department of Applied Physics, Stanford University, Stanford, CA 94305 USA; Department of Neurobiology, Stanford University, Stanford, CA 94305 USA; Advanced Photon Source, Argonne National Laboratory, Lemont, IL, 60439; Department of Psychiatry and Behavioral Sciences, Stanford University, Stanford, CA 94305 USA

## Abstract

The enteric nervous system (ENS) populates the gastrointestinal (GI) tract and controls GI function. In contrast to the central nervous system, macrostructure of the ENS has been largely overlooked. Here, we visually and computationally demonstrate that the ENS is organized in circumferential stripes that regionally differ in development and neuronal composition. This characterization provides a blueprint for future understanding of region-specific GI function and identifying ENS structural correlates of GI disorders.

## Introduction

The gastrointestinal (GI) tract is the only peripheral organ with a dedicated nervous system, which is sufficient to autonomously control GI function. Neurons of the enteric nervous system (ENS) are located within either the myenteric plexus (MP), which controls motility, or the submucosal plexus (SMP), which regulates luminal secretion and absorption. These plexuses develop from migrating neural crest cells (NCCs), which differentiate and cluster into ganglia. Disrupting ganglia formation, colonization, or neuronal tract development, such as through genetic knockout of cell adhesion genes, can have profound impacts on ENS organization and GI function^1–3^.

Particular digestive functions occur in specific GI regions, thus local structure and ENS composition may differ correspondingly to achieve these functions^4,5^. Nevertheless, unlike precisely defined pathways and structures within the brain or spinal cord^6,7^, how neurons are organized within plexuses above the level of ganglia, and how organization differs along the length of the GI tract, are poorly characterized. Here, we present a bird’s eye view of the mouse ENS, describing the previously underappreciated regional organization of the ENS, the development of this organization, and the distribution of neuronal subpopulations that comprise it.

## Results

To understand the broad layout of neurons in the ENS, we imaged large areas (up to ~50 mm^2^) of both MP and SMP in each intestinal region: duodenum, jejunum, ileum, proximal colon and distal colon. Immunolabelling neuron somas in the MP revealed a circumferential orientation of ganglia, which in turn loosely coalesced into a noncontinuous stripe pattern perpendicular to its longitudinal axis, herein referred to as neuronal stripes (Fig. 1a). In contrast, submucosal ganglia of the small intestine (SI) did not show any such arrangement, nor did they correlate with the location or organization of epithelial crypts (Fig. 1b). Ganglia of the proximal colon, on the other hand, were organized into diagonal stripes, converging directly opposite the mesenteric border and aligning with the mucosal folds of the proximal colon (Fig. 1b). Full thickness wholemount preparations did not suggest any structural interplexus relationship, and highlighted the orientation difference between the MP and SMP in the proximal colon (Fig. 1c). The stripe formations of the MP emerged as peaks in longitudinal axis signal intensity profiles of wholemount images, which also confirmed the lack of circumferential structure in the SMP (Fig. 1d). We confirmed this ENS organization using a separate imaging approach, synchrotron source x-ray tomography, performed on intact intestine that allows simultaneous visualization of multiple cell types of both MP and SMP (Fig. 1e,f). Taken together, the MP, but not the SMP, displays a striped neuronal architecture.

**Fig. 1.**
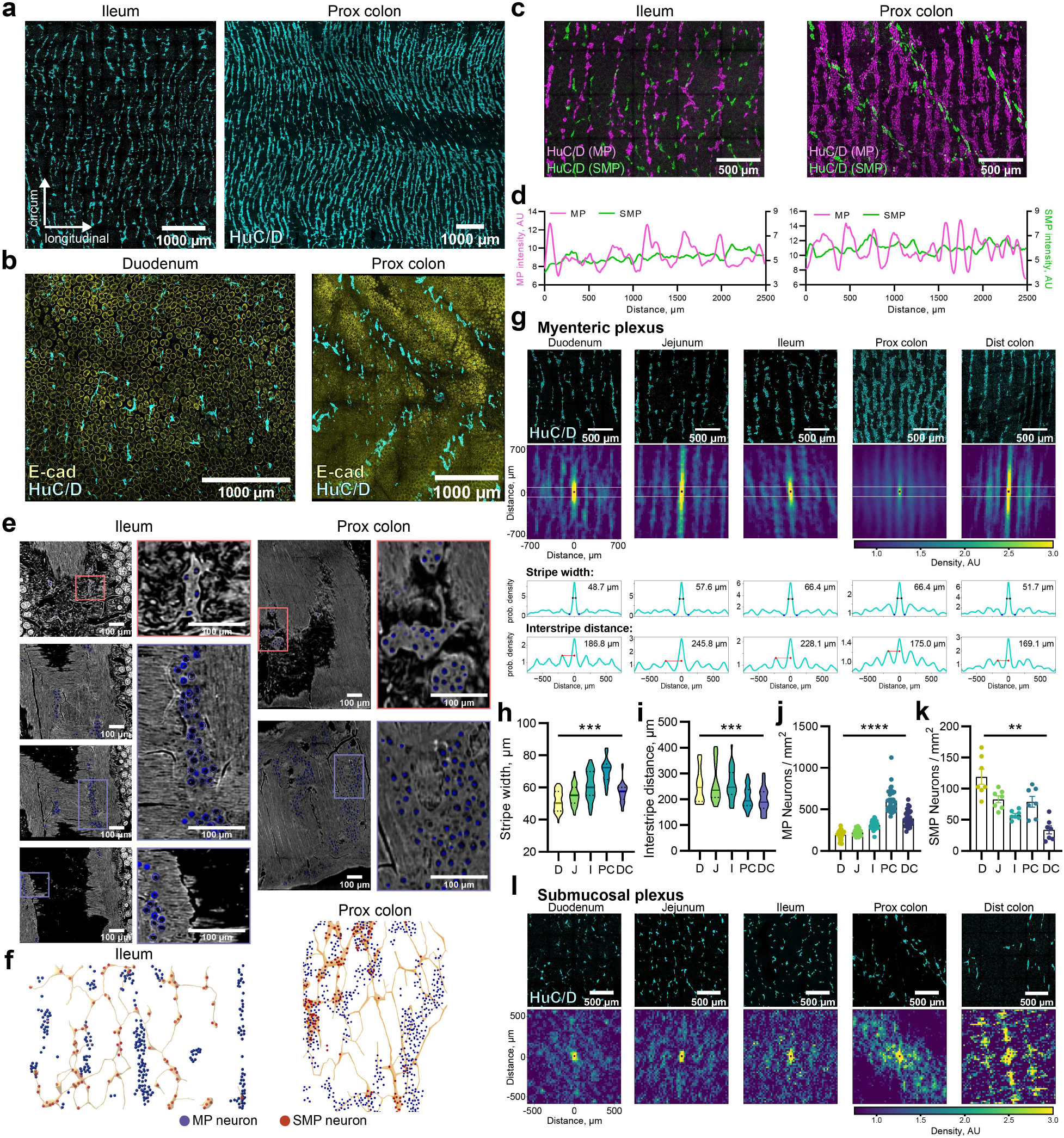
Neuronal organization in the adult ENS. **a,b,** Representative images of immunohistochemical staining of adult wholemount MP (**a**) of ileum (left) and PC (right) for neuronal label HuC/D and SMP (**b**) of duodenum (left) and PC (right) for HuC/D (cyan) and epithelial label E-cadherin (yellow). **c**, Representative images of immunohistochemical staining of full-thickness wholemount tissue of ileum (left) and proximal colon (right) for HuC/D. MP: magenta, SMP: green. **d**, Smoothed profiles of HuC/D signal intensity along the longitudinal (x-)axis of images in **c**, highlighting stripes (peaks) in MP and lack thereof in SMP. **e**, Representative x-ray tomography images of heavy metal staining of adult ileum (left) and PC (right). Colored rectangles represent locations of magnified images. Cell bodies identified based on morphology and location, annotated with blue circles. **f**, Schematic of MP (blue) and SMP (red) cell body locations from samples in **e**. Yellow shading represents the SMP. **g**, Organizational analysis of wholemount MP HuC/D immunohistochemistry of all intestinal regions (top) using conditional intensity functions (CIFs; 2^nd^ row), showing probability density of neuron locations. A value of 1 is expected density based on uniform neuron distribution. Yellow: high probability density; blue: low. Axes apply to all panels. CIFs collapsed onto 2D plot to calculate 50% stripe width (3^rd^ row, analysis of region within white lines on CIF, black dots: width measurement, blue dots: trough) and interstripe distance (bottom, includes full CIF plot, distance calculated between red dots). Inset values show given stripe width and interstripe distance values for representative samples. **h,i**, Violin plots of stripe width (**h**) and interstripe distance (**i**) from MP wholemount preparations analyzed by CIFs as in (**g**). n = 34 for all groups (**h**) and n = 21-29 (**i**). **j,k**, Neuronal density (mean ± SEM) in MP (**j**) and SMP (**k**) across intestinal regions. n = 34 (**j**) and 7 (**k**). **l**, Organizational analysis of wholemount SMP HuC/D immunohistochemistry of all intestinal regions (top) using CIFs (2^nd^ row) as in **g**. Lack of obvious stripe structure precluded further analysis. All tests one-way ANOVA, other than (**i**) which used a mixed-effects model. Pairwise comparisons not shown. \*\**p*<0.01, \*\*\**p*<0.001, \*\*\*\**p*<0.0001. Scale bars as indicated. AU, arbitrary units; D, duodenum; J, jejunum; I, ileum; PC, proximal colon; DC, distal colon; MP, myenteric plexus; SMP, submucosal plexus.

We next sought to characterize and quantify this region- and plexus-specific ENS structure. While intensity profiles (Fig. 1d) are useful visualizations, they cannot reveal any structure that is not perpendicular to the x-axis, such as in the proximal colon SMP, and they are susceptible to distortions in tissue preparation. We therefore used conditional intensity function (CIF) plots to generate spatial probability maps of neuronal locations relative to a given neuron (Fig. 1g; see Methods). The analysis revealed that an average neuron was always situated within a stripe and was flanked by higher order stripes, clearly displayed when the CIF is collapsed into a two-dimensional (2D) graph (Fig. 1g, bottom). Quantification of these graphs revealed that the proximal colon contained the thickest stripes (Fig. 1h), while the largest interstripe distances were found in the SI (Fig. 1i). This result mirrored analysis of regional neuronal density, in which the proximal colon MP was found to be the most dense, and the duodenum the least (Fig. 1j). The SMP was more sparsely populated than the MP, and, in contrast to the MP, the densest region of the SMP was the duodenum (Fig. 1k). Further, CIF analysis revealed that in the SMP only the proximal colon exhibited any macrostructure beyond ganglia organization (Fig. 1l). Therefore, the organization of enteric neurons quantitatively differs between regions and plexuses of the GI tract.

To assess how neuronal stripes arise, we visualized MP neurons in intestinal wholemount preparations from late embryonic through neonatal ages (Fig. 2a and Extended Data Fig. 1a). At embryonic day (E)14.5, when enteric neurons have populated the entire length of the mouse intestines^8^, neurons were scattered in both the SI and colon (Fig. 2a-d). Beginning at E16.5 in the jejunum and E18.5 in the distal colon, MP neurons reorganized from scattered cells into circumferential neuronal stripes, which resolved into individual ganglia at postnatal stages (Fig. 2a,b). CIF analysis confirmed the gradual emergence of neuronal stripes in the developing ENS and the different organizational timelines between the SI and colon (Fig. 2b). To quantify the progressive organization of the developing ENS, we analyzed nearest-neighbor distances and compared them to synthetically generated data with an imposed minimum nearest-neighbor distance of 10 μm, approximately the diameter of an enteric neuron (Fig. 2c). At E14.5, the dispersion of MP neurons did not differ compared to random distributions; this dispersion transitioned to highly clustered at E16.5 in the duodenum and jejunum and at early postnatal stages in the ileum and colon (Fig. 2d). Thus, neuronal stripes arise from a gradual reorganization of neurons, which occurs embryonically in the more proximal intestine and neonatally in the more distal intestine.

**Fig. 2.**
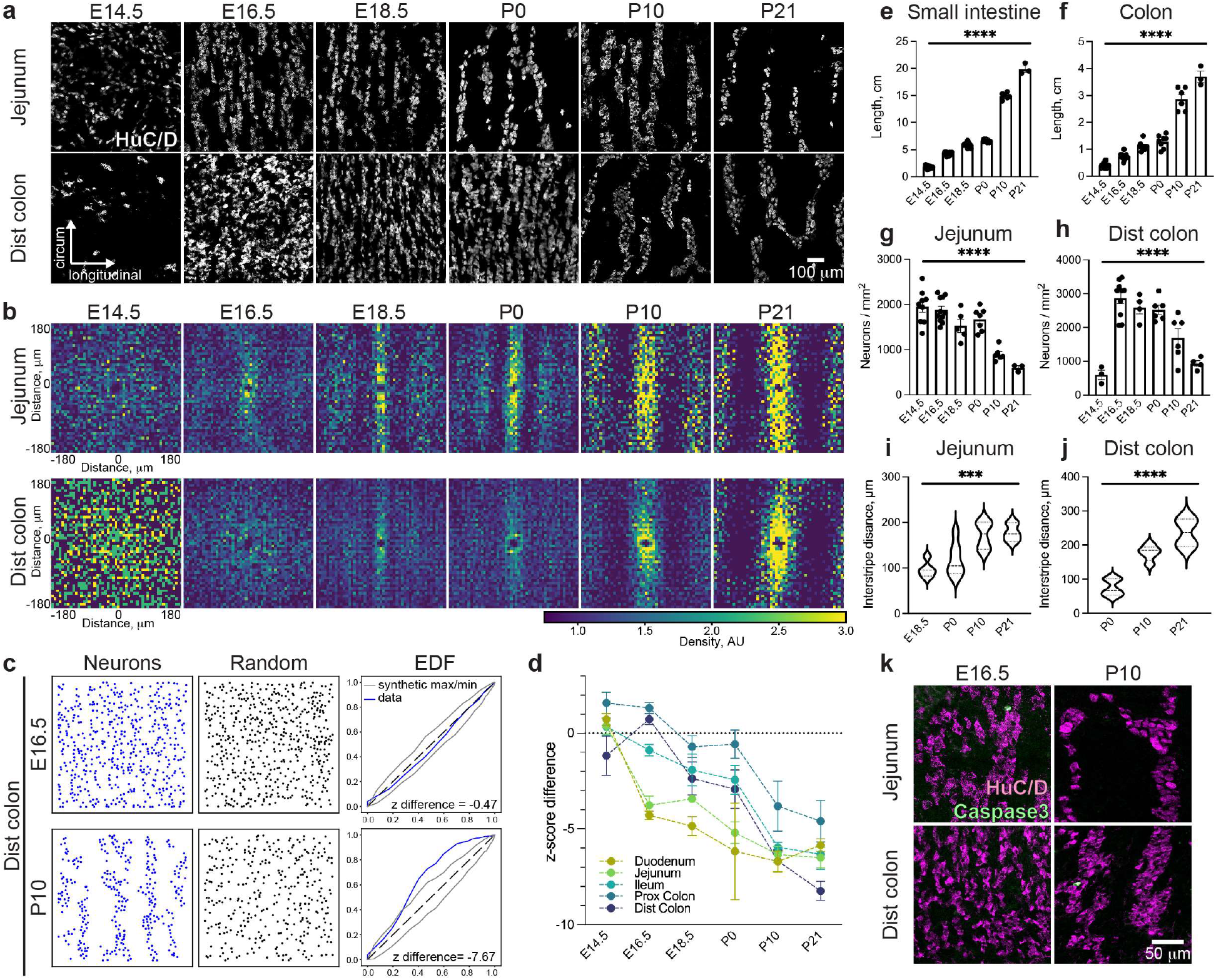
Progressive organization of the developing MP. **a**, Representative images of immunohistochemical labelling with neuronal marker HuC/D in intestinal wholemount preparations from the jejunum and distal colon MP over developmental time. Images are a subset of those in Extended Data Figure 1a. **b**, CIF plots of wholemount MP HuC/D immunohistochemistry in the jejunum and distal colon over developmental time. Yellow: high probability density; blue: low. Axes apply to all panels. **c**, 2D arrays of neurons (HuC/D+ cells) and corresponding synthetically generated data for the E16.5 and P10 distal colon. Empirical distribution function (EDF) plots for neurons (blue line) and synthetically generated maximum and minimum (gray lines). Inset values indicate z-score difference between data and mean of synthetic values for a representative sample. **d**, z-score difference between data and mean of synthetically generated values (mean ± SEM) derived from EDF plots in all regions of the intestine over developmental time. n = 2-11. **e-f**, Lengths of small intestine (**e**) and colon (**f**) (mean ± SEM). n = 3-15. **g-h**, Neuronal density (mean ± SEM) in the jejunum (**g**) and distal colon (**h**). n = 3-11. **i-j**, Violin plots of interstripe distance (mean ± SEM) in the jejunum (**i**) and distal colon (**j**) as analyzed by CIFs, n = 3-11. **k**, Representative images of immunohistochemical labelling with HuC/D (magenta) and apoptotic marker Caspase3 (green) in wholemount preparations of the jejunum and distal colon MP. All tests one-way ANOVA. Pairwise comparisons not shown. \*\*\**p*<0.001, \*\*\*\**p*<0.0001. Scale bars as indicated. AU, arbitrary units; EDF, empirical distribution function; CIF, conditional intensity function; E, embryonic day; MP, myenteric plexus; P, postnatal day.

We next assessed whether gut growth or neuronal cell death contribute to the emergence of neuronal stripes. Intestinal length increased ten-fold between E14.5 and postnatal day (P)21, which correlated with decreased neuron density and increased interstripe distance (Fig. 2e-j and Extended Data Fig. 1b-g). Expression of the apoptotic marker Caspase3 revealed sparse apoptotic neurons in the developing MP (<0.5% of HuC/D+ enteric neurons across regions and developmental time, data not shown). Collectively, these results suggest that gut growth, but not neuronal cell death, may influence organization of the developing MP.

We next sought to assess the regional distribution of neuronal subtypes, focusing on known determinants of subtype function, such as Ca2+ binding protein (CBP; Fig. 3a-c) neurotransmitter (Fig. 3d-g), and neuropeptide (Fig. 3h-k) expression^9–11^. We typically observed a greater proportion of total neurons expressing a given marker in the SI than the colon. There were particularly pronounced differences for CBPs calbindin and secretagogin (Fig. 3b,c,m,n and Extended Data Fig. 2b,c), choline acetyltransferase(ChAT)Cre-tdT (Fig. 3d,o and Extended Data Fig. 3a), and vasoactive intestinal peptide(Vip)Cre-tdT (Fig. 3j,u and Extended Data Fig. 4c), each of which labelled proportionally 2-3 fold more neurons in the SI than the colon. Tachykinin Precursor 1(Tac1)Cre-tdT showed a similarly diverse expression pattern across regions, albeit with some instances of mosaicism in the colon, where substance P immunoreactivity was observed in the absence of the tdTomato reporter (Fig. 3h,s and Extended Data Fig. 4a,e,f). The proportion of proenkephalin(Penk)Cre-tdT-expressing neurons gradually decreased along the length of the GI tract (Fig. 3i,t and Extended Data Fig. 4b,h,i). Both PenkCre-tdT and Tac1Cre-tdT also displayed some non-neuronal tdTomato expression (Extended Data Fig. 4g,j). Interestingly, somatostatin was the only marker more prevalent in the colon than in the SI (Fig. 3k,v and Extended Data Fig. 4d). It remains unclear which neuronal subtypes predominate in the distal colon.

**Fig. 3.**
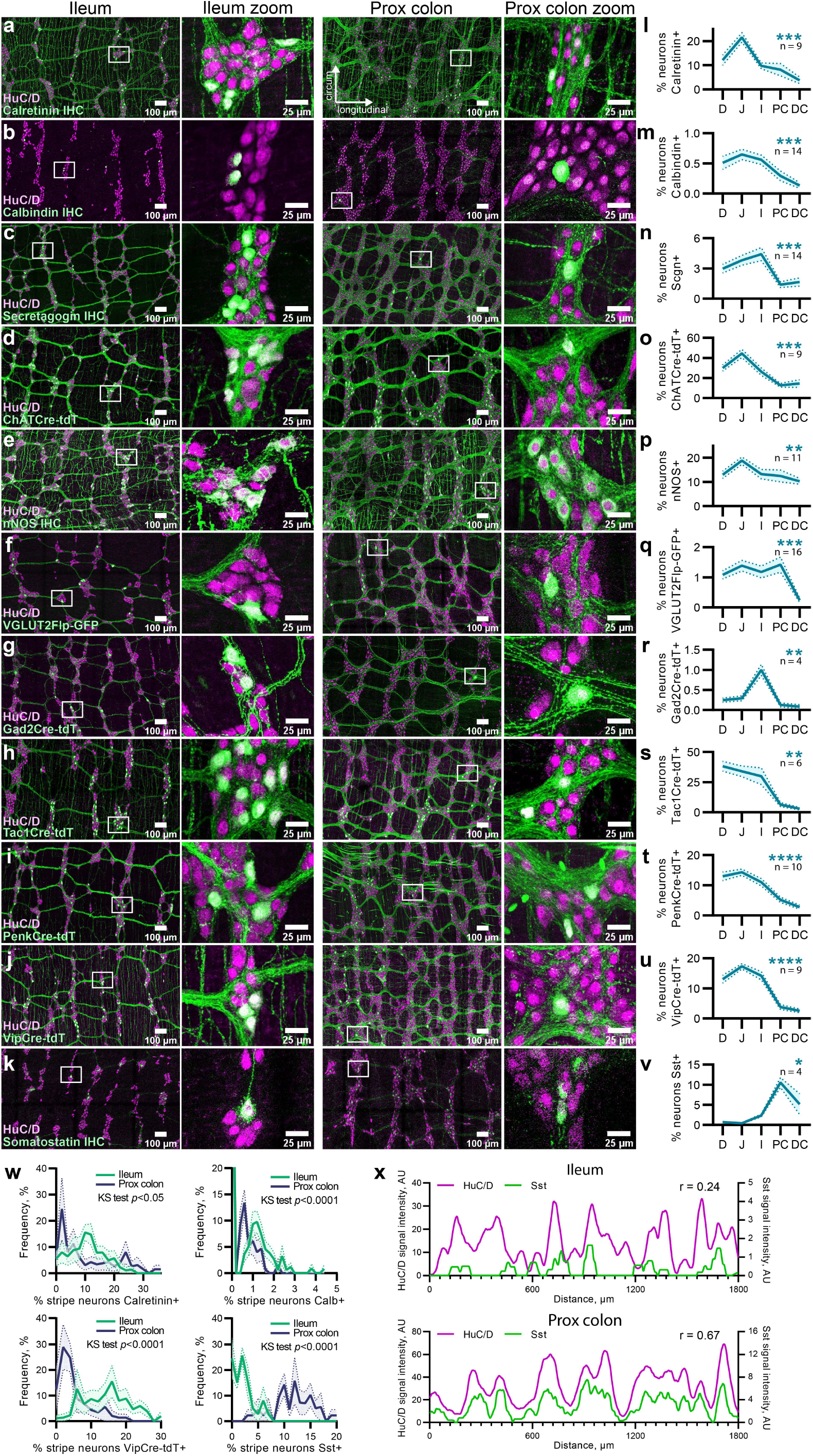
Regional distribution and organization of neuronal subtypes. **a-k,** Representative images of adult wholemount MPs showing immunohistochemical labels or genetically encoded reporters for calcium binding proteins (**a-c**), neurotransmitters (**d-g**) and neuropeptides (**h-k**) alongside neuronal label HuC/D in the ileum (left) and PC (right). Higher magnification image locations indicated by white boxes. Scale bars as indicated. **l-v**, Proportion of total HuC/D neurons (mean ± SEM) positive for each neuronal marker across intestinal regions as in **a-k**. n as indicated in graph. All tests one-way ANOVA. Pairwise comparisons not shown. \**p*<0.05, \*\**p*<0.01, \*\*\**p*<0.001, \*\*\*\**p*<0.0001. **w**, Frequency distribution plots (mean ± SEM) of the proportion of neurons in a neuronal stripe positive for calretinin (6-12 stripes per region, 8 mice), calbindin (6-14 stripes, 14 mice), VipCre-tdT (9-14 stripes, 8 mice) and somatostatin (9-11 stripes, 4 mice). **w**, Representative smoothed profiles of HuC/D and somatostatin signal intensity along the longitudinal axis in ileum (top) and PC (bottom). r value indicates Pearson r correlation between the two profiles. Calb, calbindin; ChAT, choline acetyltransferase; Cre, Cre recombinase; Flp, Flp recombinase; Gad2, glutamate decarboxylase 2; GFP, green fluorescent protein; IHC, immunohistochemistry; KS, Kolmogorov-Smirnov; nNOS, neuronal nitric oxide synthase; Penk, proenkephalin; Scgn, secretagogin; Sst, somatostatin; Tac1, Tachykinin Precursor 1; tdT, tdTomato; VGLUT2, vesicular glutamate transporter 2; Vip, vasoactive intestinal peptide.

We also assessed whether marker expression differed between regions within the SI or colon. Significantly higher expression of calretinin (Fig. 3a,l and Extended Data Fig. 2a), ChATCre-tdT (Fig. 3o) and neuronal nitric oxide synthase (nNOS; Fig. 3e,p and Extended Data Fig. 3b) was found in the jejunum than in any other region. Vesicular glutamate transporter 2(VGLUT2)Flp-GFP was significantly lower in the distal colon compared to all other regions (Fig. 3f,q and Extended Data Fig. 3c,f), and glutamate decarboxylase 2(Gad2)Cre-tdT expression was ~4 fold higher in the ileum (Fig. 3g,r and Extended Data Fig. 3d). Thus, distribution of key neuronal subtypes differs considerably both between and within the SI and colon.

The low expression of markers in the colon was surprising, not least that of ChATCre-tdT, as cholinergic communication is widely regarded as a primary form of enteric neurotransmission. Despite this, mean expression of ChATCre-tdT did not exceed 50% of neurons in any region, and was less than 20% in both regions of the colon (Fig. 3o). This distribution was supported by strong colocalization between ChATCre-tdT and vesicular acetylcholine transporter (vAChT) IHC (Extended Data Fig. 3e). Further, even concomitant visualization of three major enteric neurotransmitters by nNOS immunostaining of VGLUT2Flp-GFP-ChATCre-tdT tissue left a majority of HuC/D-positive neurons lacking any of the three markers (Extended Data Fig. 3g-i). Therefore, the neurotransmitter used by ChATCre-tdT/nNOS/VGLUT2Flp-GFP-negative neurons remains unknown.

Despite distribution differences in cell body localizations, staining of neuron fibers tended to be qualitatively similar between regions, for example ChATCre-tdT fibers were observed in circular muscle in both SI and colon. The exception to this was secretagogin. In the SI, secretagogin fibers were restricted to interganglionic tracts, while they additionally innervated longitudinal and circular muscle in the proximal and distal colon, respectively (Fig. 3c and Extended Data Fig. 2c). This may suggest a different, as yet uncharacterized, function for secretagogin neurons, dependent on intestinal region.

We next examined neuronal subtype representation across myenteric neuronal stripes. We selected 4 markers for this analysis, covering a range of different cell types: motor neurons (VipCre-tdT and calretinin), sensory neurons (calbindin) and interneurons (somatostatin). At the level of enteric neuronal stripes, the proportion of neurons positive for a given marker broadly reflected that of the entire region, and these distributions shifted appropriately between regions (Fig. 3w). For instance, VipCre-tdT is in ~15% of ileum neurons overall (Fig. 3u), and each stripe was found to have 0-30% VipCre-tdT neurons, with a peak at 15% (Fig. 3w). Subtypes appear to be well distributed across stripes, with few stripes containing zero neurons of a given subtype unless that marker is expressed in very few cells, such as calbindin (Fig. 3b,w). This is further illustrated when comparing longitudinal axis intensity profiles of HuC/D signal with that of a given subtype (Fig. 3x and Extended Data Fig. 5). Only at very low levels did the correlation between profiles vanish, as seen in the somatostatin ileum profile compared with the colon profile (Fig. 3x). Indeed, the strength of this correlation greatly increased over a small range when plotted against the proportion of neurons positive for a given marker (Extended Data Fig. 5d). This suggests that, even with a restricted area of analysis (1800 μm^2^), neuron subtype prevalence can be low (2-3%), yet its spatial distribution within a region can reflect that of HuC/D, providing evidence for an approximately even distribution of a given subtype across enteric neuronal stripes.

## Discussion

By combining large-scale image analysis with computational methods over multiple regions and ages of the mouse intestine, we demonstrate that the mouse MP possesses a gross cytoarchitecture of circumferential neuronal stripes that differ by region in their organization, development, and neuronal composition.

Differences in ENS organization and composition may contribute to the unique functions within each GI region. Single-cell RNA sequencing has revealed differential distribution of functional neuronal classes throughout the GI tract^12,13^, observations confirmed by our results and extended to the level of neuronal stripes. This diversity differs more region-to-region than among neuronal stripes of a given region, which tend to possess similar neuronal compositions to their neighbors.

The role of individual stripes remains to be elucidated. Clonally related ENS cells inhabit overlapping domains and exhibit coordinated activity^14^; whether these functional units map onto the macrostructure of neuronal stripes has yet to be explored. Further, stripes may serve to control rings of circular muscle, which possess the same orientation as neuronal stripes, or to coordinate longitudinal interneuron projections oriented in parallel^15^.

Another outstanding question is the mechanism that underlies the development of neuronal stripes. The spatiotemporal differences we observe in neuronal stripe development may reflect timing of NCC colonization. Development of circular muscle may also influence this patterning. The circular muscle differentiates around the time of neuronal stripe emergence^16,17^, and inhibition of circular muscle contractions changes the anisotropy of the embryonic chick ENS^18^. Such mechanistic insights may improve methods for generating intestinal organoids, examples of which have two ENS plexuses but lack patterning along the longitudinal axis^19,20^.

Our characterization of regional enteric organization and composition can serve as a blueprint to understand how local enteric neurocircuitry underlies region-specific motility patterns and function. Further, assessing ENS structure may help identify anatomical changes in human GI diseases with no known correlates within the ENS, such as intestinal pseudo-obstruction^21^. Comprehensive analyses of central nervous system structure have served as platforms to dissect basic physiology and disease, and our work represents a parallel step toward an intricate understanding of the ENS.

## Methods

### Animals

Animals were group-housed, and all procedures conformed to the National Institutes of Health Guidelines for the Care and Use of Laboratory Animals and were approved by the Stanford University Administrative Panel on Laboratory Animal Care or the University of Chicago Institutional Animal Use and Care Committee. For embryonic experiments, female mice were visually inspected for the presence of a vaginal plug every morning, and plugged females were separated from breeding cages.

X-ray tomography data was obtained from 13-15 week-old female C57BL/6J mice (Jackson, #000664). All experiments on the development of enteric neuron organization were performed on C57BL/6J mice (Jackson, #000664) at the ages indicated. Experiments assessing neuronal subtype distribution used the following recombinase lines: ChATCre (B6.129S-*Chat^tm1(cre)Lowl^*/MwarJ (Jackson, #031661)), VGLUT2Flp (B6;129S-*Slc17a6^tm1.1(flpo)Hze^*/J (Jackson, #030212)), Gad2Cre (STOCK *Gad2^tm2(cre)Zjh^*/J (Jackson, #010802)), VipCre (STOCK *VIP^tm1(cre)Zjh^*/J (Jackson, #010908)), Tac1Cre (B6;129S-*Tac1^tm1.1(cre)Hze^*/J (Jackson, #021877; kind gift from William Giardino)), and PenkCre (B6;129S-*Penk^tm2(cre)Hze^*/J (Jackson, #025112)). These recombinase lines were then crossed with either GFP (STOCK *Gt(ROSA)26Sor^tm1.2(CAG-EGFP)Fsh^*/Mmjax (Jackson, #32038-JAX)) or tdTomato (B6;129S6-*Gt(ROSA)26Sor^tm14(CAG-tdTomato)Hze^*/J (Ai14; Jackson, #007908)) lines as appropriate. Investigations assessing general neuronal structure of the ENS collated data from all of the above lines, from both sexes, aged 2-6 months.

### Marker selection

For experiments assessing neuronal subtype distribution, we selected a variety of markers (via either genetically encoded reporters, above, or antibodies (Table1)) covering calcium binding proteins, neurotransmitters and neuropeptides, which are known determinants of enteric subtype function or which delineate functionally relevant populations^13,22^. Furthermore, all labels chosen marked the neuron soma, which facilitated automated counting of large areas of tissue. Calretinin, ChAT, and Tac1 are all known to mark excitatory motor neurons. Penk also marks excitatory motor populations, in addition to some interneurons. Somatostatin has been suggested to delineate a population of interneurons^23^. Inhibitory motor neurons were labelled by VIP and nNOS. Calbindin is suggested to mark sensory neurons^24^. The functions of VGLUT2 and Gad2 neurons are not known, but were included as they are well-known neurotransmitter markers in other areas of the nervous system and had been identified in the ENS by recent scRNAseq studies as belonging to specific subpopulations^13,25^. Secretagogin, a calcium binding protein first identified in the pancreas^26,27^, was similarly identified recently, and has not yet been described in the ENS.

**Table 1:** Primary antibodies used for immunohistochemistry

| **Antibody** | **Host** | **Dilution** | **Source** | **Catalogue No.** | **RRID** |
| --- | --- | --- | --- | --- | --- |
| **Calbindin** | Mouse | 1:1000 | Swant | CB300 | AB_10000347 |
| **Calretinin** | Rabbit | 1:2000 | Millipore | AB5054 | AB_2068506 |
| **Caspase3** | Rabbit | 1:500 | Abcam | AB2302 | AB_302962 |
| **E-cadherin** | Rat | 1:2000 | ThermoFisher | 13-1900 | AB_2533005 |
| **GFP** | Sheep | 1:1000 | Biogenesis | 4745-1051 | AB_619712 |
| **HuC/D** | Human | 1:75000 | Mayo Clinic | King et al., 1999<sup>28</sup> |  |
| **Met-Enkephalin** | Rabbit | 1:1000 | Immunostar | 20065 | AB_572250 |
| **nNOS** | Sheep | 1:1000 | Millipore | AB1529 | AB_90743 |
| **nNOS** | Rabbit | 1:2000 | Sigma-Aldrich | N7280 | AB_260796 |
| **Secretagogen** | Chicken | 1:3000 | EnCor Bio | CPCA-SCGN | AB_2744521 |
| **Somatostatin** | Rat | 1:500 | Millipore | MAB354 | AB_2255365 |
| **Substance P** | Rabbit | 1:4000 | Immunostar | 20064 | AB_572266 |
| **vAChT** | Goat | 1:2000 | Chemicon | AB1578 | AB_10000324 |
| **VGLUT2** | Guinea pig | 1:3000 | Synaptic Systems | 135 404 | AB_887884 |

### Immunohistochemistry

For investigations into the structure of the adult ENS, the small intestine and colon were dissected out from 2-6 months-old mice culled by CO_2_ and cervical dislocation. Intestines were flushed of fecal contents using cold PBS before cutting the proximal, middle and distal 2-3 cm of the small intestine to isolate the duodenum, jejunum and ileum. The colon was cut in two, and all segments were placed in cold PBS on ice. A fire-polished glass rod 2 mm in diameter was placed through each segment, and the mesentery removed. For full-thickness wholemount preparations, tissue was fixed on the glass rod in 4% PFA for 90 minutes at 4°C with shaking. For adult single-plexus wholemount preparations, the muscularis (both muscle layers and MP) was peeled away using cotton buds, starting at the mesenteric border. For neonatal ages (postnatal days 0, 10, 15, and 21), intestines were cut open along the mesenteric border, pinned mucosa-down on Sylgard 170 in ice-cold PBS, and the muscularis was peeled away with fine forceps. Muscularis tissue was stored in ice-cold PBS and then pinned onto Sylgard 170 in a flat sheet in 35 mm glass dishes using insect pins. In some cases, the underlying mucosa containing the SMP was also kept and pinned. Tissue was immediately fixed as described above. Residual PFA was removed via PBS washes before proceeding directly to immunostaining, or stored in PBS containing 0.1% sodium azide for up to 3 weeks at 4°C.

For embryonic experiments, pregnant mice were culled by CO_2_ and cervical dislocation. The uterus was removed, and embryos were placed in ice-cold PBS. The intestines were dissected from each embryo, and the mesentery was carefully removed. To measure intestinal length, the cleaned intestine was laid adjacent to a ruler. In a Sylgard 170 plate, a pin was placed in the stomach and anus to keep the intestines taut and straight. Tissue was fixed in 4% PFA for 90 minutes at 4°C with shaking.

For immunohistochemistry, large intact pieces of tissue were cut into smaller pieces and transferred to WHO microtitration trays (International Scientific Supplies) containing PBS. Tissue was then put into PBT (PBS, 1% BSA, 0.1% Triton X-100) containing the primary antibodies (see Table 1) overnight at 4°C with shaking. For mouse antibodies, tissue was pre-treated with Affinipure FAb Fragment Donkey Anti-mouse (Jackson 715-007-003) diluted 1:50 in PBT, and in 5% normal goat serum in PBT, for 2 h each prior to primary antibody incubation^13^. The following day, tissue was washed 3 times in PBT for 30 minutes each before transferring to PBT containing secondary antibodies (Table 2) for 2 h at room temperature with shaking. Tissue was washed twice in PBT and twice in PBS, then mounted onto slides, rinsed in ddH_2_O and coverslipped using Fluoromount-G (Southern Biotech).

**Table 2:** Secondary antibodies used for immunohistochemistry

| **Target** | **Fluorophore** | **Host** | **Dilution** | **Source** | **Catalogue No.** | **RRID** |
| --- | --- | --- | --- | --- | --- | --- |
| **Chicken** | Cy3 | Donkey | 1:500 | Jackson Immuno Research | 703-165-155 | AB_2340363 |
| **Chicken** | AF 647 | Donkey | 1:500 | Jackson Immuno Research | 703-605-155 | AB_2340379 |
| **Guinea pig** | AF 647 | Donkey | 1:500 | Jackson Immuno Research | 706-605-148 | AB_2340476 |
| **Goat** | AF 647 | Donkey | 1:500 | Jackson Immuno Research | 705-605-003 | AB_2340436 |
| **Human** | AF 488 | Donkey | 1:1000 | Jackson Immuno Research | 709-545-149 | AB_2340566 |
| **Human** | AF 405 | Donkey | 1:500 | Jackson Immuno Research | 709-475-149 | AB_2340553 |
| **Human** | AF 647 | Donkey | 1:500 | Jackson Immuno Research | 709-605-149 | AB_2340578 |
| **Mouse** | Cy3 | Goat | 1:500 | Jackson Immuno Research | 115-165-205 | AB_2338694 |
| **Mouse** | AF 647 | Goat | 1:500 | Jackson Immuno Research | 115-605-205 | AB_2338916 |
| **Rabbit** | AF 488 | Donkey | 1:500 | Invitrogen | A21206 | AB_2535792 |
| **Rabbit** | Cy5 | Donkey | 1:500 | Jackson Immuno Research | 711-175-152 | AB_2340607 |
| **Rabbit** | Cy3 | Donkey | 1:800 | Jackson Immuno Research | 711-165-152 | AB_2307443 |
| **Rat** | AF 488 | Donkey | 1:500 | Invitrogen | A21208 | AB_141709 |
| **Rat** | Cy3 | Donkey | 1:500 | Jackson Immuno Research | 712-165-153 | AB_2340667 |
| **Sheep** | AF 488 | Donkey | 1:500 | Invitrogen | A11015 | AB_141362 |
| **Sheep** | Cy5 | Donkey | 1:500 | Jackson Immuno Research | 713-175-147 | AB_2340730 |

For embryonic experiments, immunohistochemistry protocol was done as in adult with the following changes. Intact intestines were placed in WHO microtitration trays (International Scientific Supplies) and treated as above. After immunohistochemistry, fixed intestines were cannulated with a cleaning wire for 33-gauge needles (Hamilton) and cut along the wire. Embryonic intestines were mounted full-thickness.

### X-ray tomography

Mice were deeply anesthetized using pentobarbital (60 mg/kg intraperitoneal) to be non-responsive to toe pinch and transcardially perfused with 10 ml of 0.1 M sodium cacodylate buffer (Electron Microscopy Sciences (EMS)) to flush the vasculature followed by buffered fixative made of 2.5% glutaraldehyde (EMS), 2% paraformaldehyde (EMS) in 0.1 M sodium cacodylate buffer (EMS). The intestines were removed, and the mesentery was gently severed. The intestines were divided into segments (duodenum, jejunum, ileum, proximal and distal colon). Ileum and proximal colon were dissected out and gently flushed with 0.1 M sodium cacodylate buffer (EMS). The dissected segments were post-fixed overnight at 4°C in the fixative described above. After post-fixation, the intestines were cut open along the mesentery and dissected into smaller pieces before staining with heavy metals as described by Hua et al^29^. Briefly, the tissues were extensively washed in 0.1 M sodium cacodylate buffer (EMS). This was followed by sequential staining with 2% buffered osmium tetroxide (EMS), 2.5% potassium ferrocyanide (Sigma-Aldrich) with no rinse in between followed by pyrogallol (Sigma-Aldrich), unbuffered 2% osmium tetroxide (EMS), 1% uranyl acetate (EMS), and 0.66% aspartic acid buffered lead (II) nitrate (Sigma-Aldrich) with extensive rinses in between each of the steps. The stained tissues were dehydrated in graded ethanol, propylene oxide and infiltrated with epon resin (EMS). The tissues were finally embedded in fresh epon and cured in an oven at 60°C for 48 h^29,30^.

#### Synchrotron imaging

The epon embedded samples were imaged at 32-ID beamline at the Advanced Photon Source (APS) in Argonne National Laboratory. The sample was placed on a rotation stage and projection images were acquired at 600 nm/pixel resolution as the sample was rotated over 180 degrees^31,32^. The acquired images were 3D-reconstructed using the TomoPy toolbox^33^.

#### Data Analysis

The 3D-reconstructed data were manually annotated using Knossos, an open-source software (https://knossos.app/)^34^. We identified the myenteric ganglia in between the circular and longitudinal muscle layers and the submucosal ganglia in the submucosal space. All cell bodies were identified based on the circular outline and the presence of a nucleus within the cell.

### Image acquisition

Images were acquired using a 20x (NA 0.75) oil objective on a Leica SP8 confocal microscope. Large tiled images were acquired and stitched using the Navigator mode within LASX. Z-stacks were acquired with 3 μm between each focal plane for adult tissue and with 2 μm between each focal plane for embryonic and neonatal tissue.

### Image Analysis

#### Cell counting

Image analysis was primarily performed using ImageJ/FIJI (NIH, Bethesda, MD). For neuronal density analysis, HuC/D images (maximum intensity projections for adults, single plane for perinatal) were blurred before thresholding and watershedding. Cells were counted using the Analyze Particles function, and density was calculated based on area of tissue measured in FIJI. To count neuronal subtypes, the Image Calculator function was used to combine thresholded HuC/D stacks (prior to maximum projection) with raw image stacks of a defined cell type. The result of this calculation was then maximally projected and counted as above.

#### Conditional intensity function (CIF) analysis

To visualize and evaluate spatial patterns, we calculated the conditional intensity function (CIF), which generates a spatial density map of neuron locations relative to a given neuron. Square images measuring ~1800×1800 μm for adult, ~400×400 μm for embryonic and neonatal tissue, respectively, were processed in FIJI as described above, and the XY coordinates of each neuron were obtained using Analyze Particles. We empirically estimated the CIF for a given sample by iterating over all neurons and calculating the number of neurons in a 2D grid around that neuron. Total image area was normalized to 1, divided into 100 bins per unit length. The 2D grid’s width and height were both 0.7 for adult data, while they were 0.8 and 0.5, respectively, for embryonic/neonatal data. Density values were normalized to expected density based on a uniform distribution of neurons, given a value of 1. We excluded the center grid point from the resulting CIF plot, which included data from the neuron used for conditioning.

We then transformed the 2D grid into a one-dimensional line by averaging along the y-axis, using either the full y-axis (for interstripe distance calculations) or a smaller proportion (for stripe width calculations). For adult data, this smaller proportion was a length of 0.1 relative image length above and below the center, and for developmental data, we used a value of 0.2. For developmental data, which contained far fewer neurons, we smoothed with a Gaussian of 20 μm standard deviation. We then identified the first minima and first peak next to the center. Stripe width was taken as the width at the half height from minimum to center peak. Interstripe distance was taken as the distance between the left and center peaks. For interstripe distance analysis, samples in which secondary stripes could not be unambiguously identified were not included. All analyses were implemented in Python.

#### Nearest-neighbor and Empirical distribution function (EDF) analysis

To assess enteric neuron organization across development, we tested each of the samples for deviation from a hypothesis of random neuron positions, known as complete spatial randomness. Our analysis was based on nearest neighbor distances. The distribution of nearest neighbor distances under complete spatial randomness can be calculated^35^. Specifically, the mean of the distribution and its variance can be approximated by a normal distribution^35^. Embryonic and neonatal images were processed as above to generate XY coordinates, and then for each sample we calculated the mean of the data and extracted a z-score for the deviation of the data from the expected value. In addition, to better visualize deviations, we generated synthetic samples under the assumption of complete spatial randomness. We then plotted the empirical distribution function for the data as well as an envelope defined by the maximum and minimum of the empirical distribution function over 500 synthetic samples.

We observed that some deviation from randomness was driven mostly by the fact that the synthetically generated samples exhibited overlap in space while our data, taken from a single-plane image, did not overlap. To make the analysis more robust to this property, we generated synthetic samples with a minimum distance imposed. We chose this minimum distance as 10 μm, approximately the average diameter of an enteric neuron. We then generated 500 random samples and calculated their z-score and defined a metric that is the difference between the z-score of the data and the mean of the z-score of the synthetic samples. This allowed better identification of differences from other spatial properties (e.g., stripes). All analyses were implemented in Python.

#### Longitudinal axis signal profile analysis

Profiles of HuC/D and neuronal subtype signal intensity along the longitudinal tissue axis were generated in FIJI using the Plot Profile function, once the image had been adequately aligned such that the longitudinal axis of the tissue matched the x-axis. These profiles were then exported to Prism 9 (GraphPad) for correlation analysis (see Statistical analysis). After analysis, profiles were smoothed in Prism 9 by averaging 12.5 μm either side of a given point to aid in visualizing peaks in intensity. To calculate the number of HuC/D neurons and neurons of a given subtype present in stripes within an image, smoothed profiles were first exported to Microsoft Excel and peaks in the data were automatically identified. The locations of these peaks were then exported to FIJI, which created a box measuring the full y-axis and 50 μm either side of each peak location. This box would constitute a stripe, and Analyze Particles was used on thresholded images (as above) to count the number of objects within each stripe. This data was converted to frequency distributions in Prism 9.

### Statistical analysis

Statistical tests and graphical representation of data were performed using Prism 9 software (GraphPad). Statistical comparisons were performed using one-way ANOVA to assess if age or intestinal region were significant factors for a variety of measurements (neuron density, intestine length etc.); repeated measures one-way ANOVAs were used for intestinal region. Tukey’s or Dunnett’s correction for multiple comparisons was used where appropriate to determine significant differences between individual regions or ages. Full details of these statistical tests can be found in Supplementary Table 1. Correlation was determined using Pearson’s correlation coefficient. The Kolmogorov-Smirnov (KS) test was used to compare the frequency distributions of neuron subtypes as a proportion of neurons in neuronal stripes.

### Data availability

The data that support the findings of this study are available upon reasonable request.

### Code availability

The code used to analyze the data in this study can be found at https://github.com/druckmann-lab/EntericNervousSystemAnalysis.

## Acknowledgements

We thank members of the Kaltschmidt lab for experimental advice and discussions, and Julieta Gomez-Frittelli and Subhamoy Das for feedback on the manuscript. We thank Dr. Vanda A. Lennon (Mayo Clinic) for the HuC/D primary antibody. This work was supported by an EMBO Fellowship ALTF 180-2019 (R.H.), a Stanford Medical Scientist Training Program grant T32 GM007365-44 (L.B.D.), a grant from The Leona M. and Harry B. Helmsley Charitable Trust and NIDDK P30DK042086 (V.S, N.K.), the Alfred P. Sloan Foundation, the McKnight Foundation and NIH R01EB028171 (S.D.), and the Wu Tsai Neurosciences Institute, the Stanford University Department of Neurosurgery and a research grant from The Shurl and Kay Curci Foundation (J.A.K.).

## Author Contributions

R.H. designed, performed and analyzed immunohistochemistry experiments relating to adult neuron organization and neuronal subtypes, and co-wrote the manuscript. L.B.D. designed, performed and analyzed immunohistochemistry experiments relating to development, and co-wrote the manuscript. V.S. designed, performed and analyzed x-ray tomography experiments. Z.W. and S.D. wrote the code for data analysis. V.D.A. helped collect the x-ray tomography dataset. N.K. supervised and designed x-ray tomography experiments. J.A.K. supervised and designed immunohistochemistry experiments and co-wrote the manuscript.

