## Supplementary Fig. 1 for "Regional cytoarchitecture of the adult and developing mouse enteric nervous system"

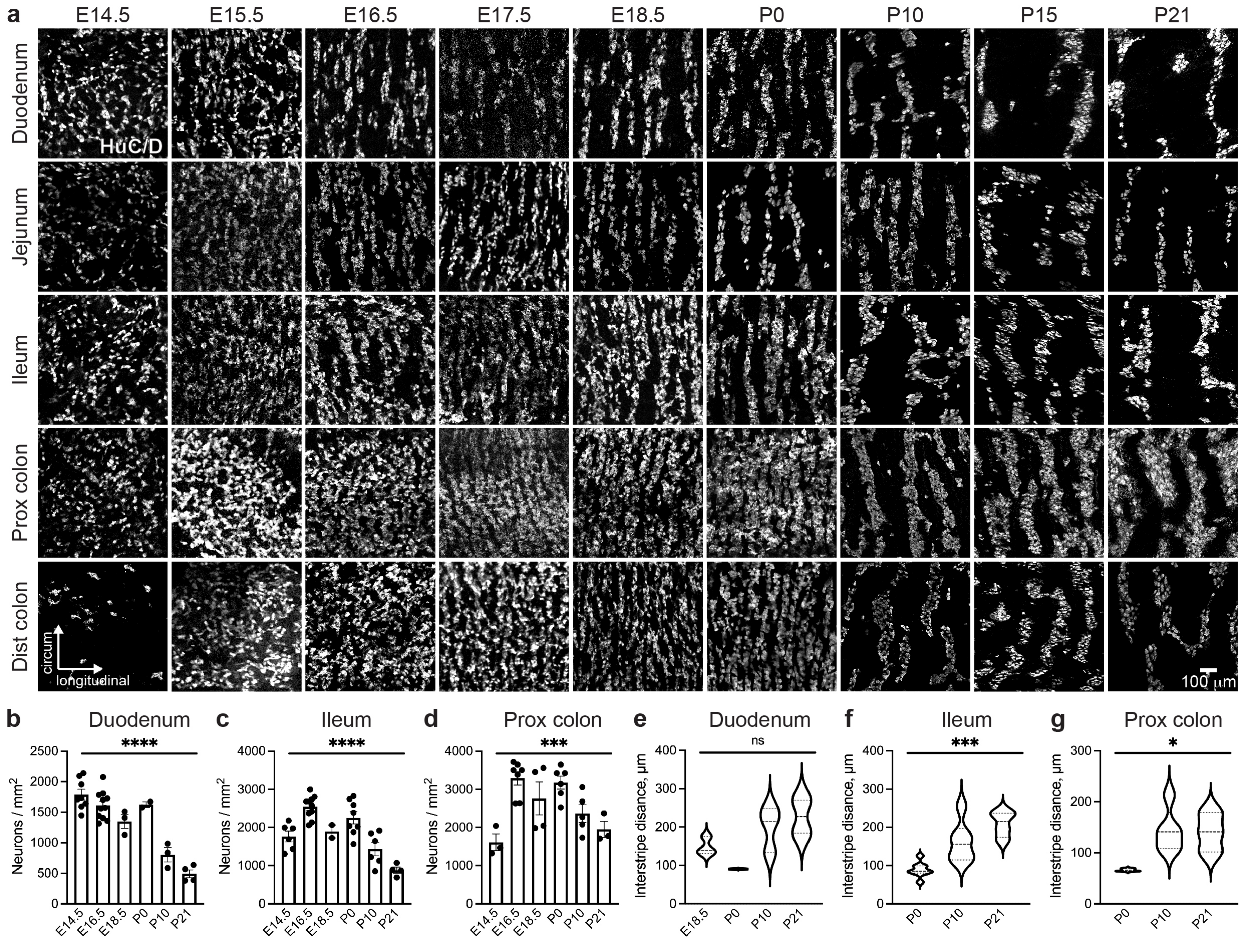


**Supplementary Fig. 1 (Supporting Figure 2) | Progressive ENS organization occurs across all regions of the developing intestine. a**, Representative images of immunohistochemical labelling with neuronal marker HuC/D in intestinal wholemount preparations from the duodenum, jejunum, ileum, proximal colon, and distal colon MP over developmental time. **b-d**, Neuronal density (mean ± SEM) in the duodenum (**b**), ileum (**c**), and proximal colon (**d**). **e-g**, Violin plots of interstripe distance in the duodenum (**e**), ileum (**f**), and proximal colon (**g**) as analyzed by CIF. n = 2-11. All tests one-way ANOVA. Pairwise comparisons not shown. *$p$<0.05, **$p$<0.01, ***$p$<0.001, ****$p$<0.0001. Scale bar as indicated. CIF, conditional intensity function; MP, myenteric plexus.
