## Supplementary Fig. 2 for "Regional cytoarchitecture of the adult and developing mouse enteric nervous system"

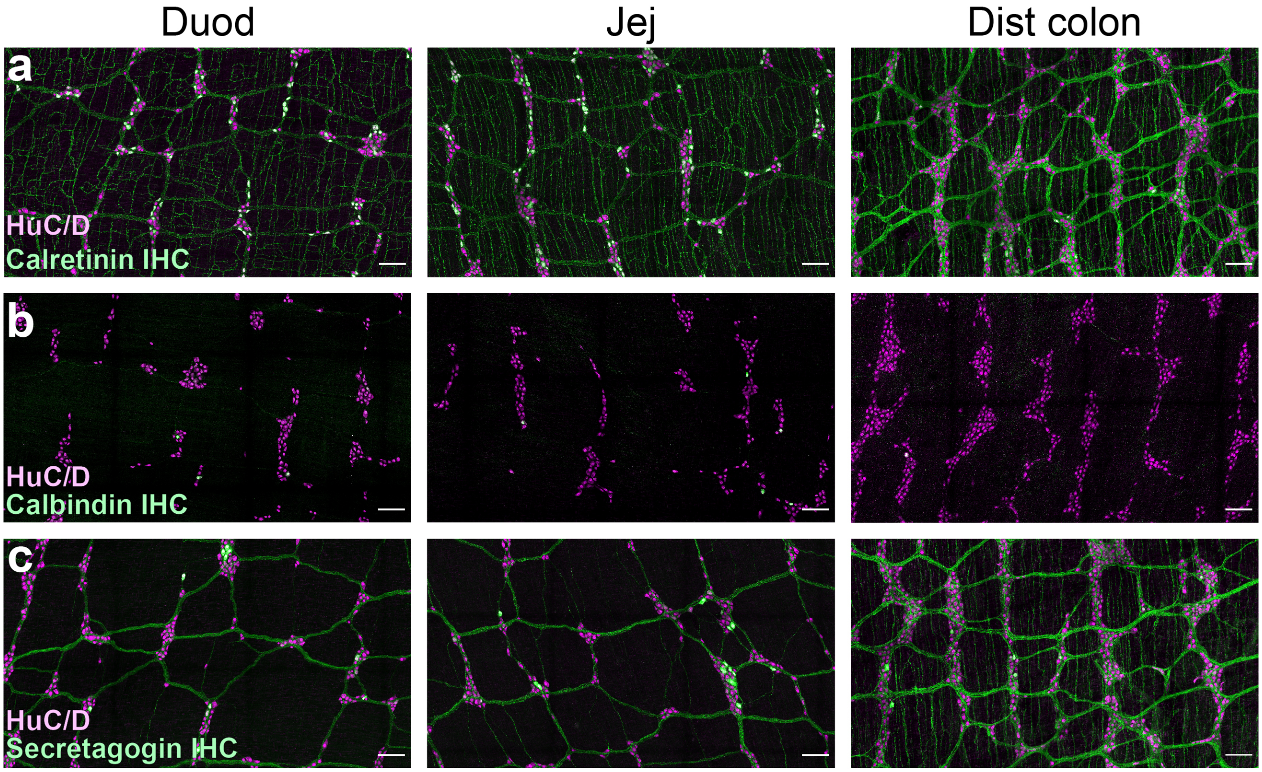


**Supplementary Fig. 2 (Supporting Figure 3) | Calcium binding protein distribution in the ENS. a-c**, Representative images of immunohistochemical labelling of adult wholemount MP duodenum, jejunum and DC for HuC/D (magenta) and calretinin (**a**), calbindin (**b**) or secretagogin (**c**) (green). Scale bars 100 µm. DC, distal colon; IHC, immunohistochemistry; MP, myenteric plexus.
