## Supplementary Fig. 3 for "Regional cytoarchitecture of the adult and developing mouse enteric nervous system"

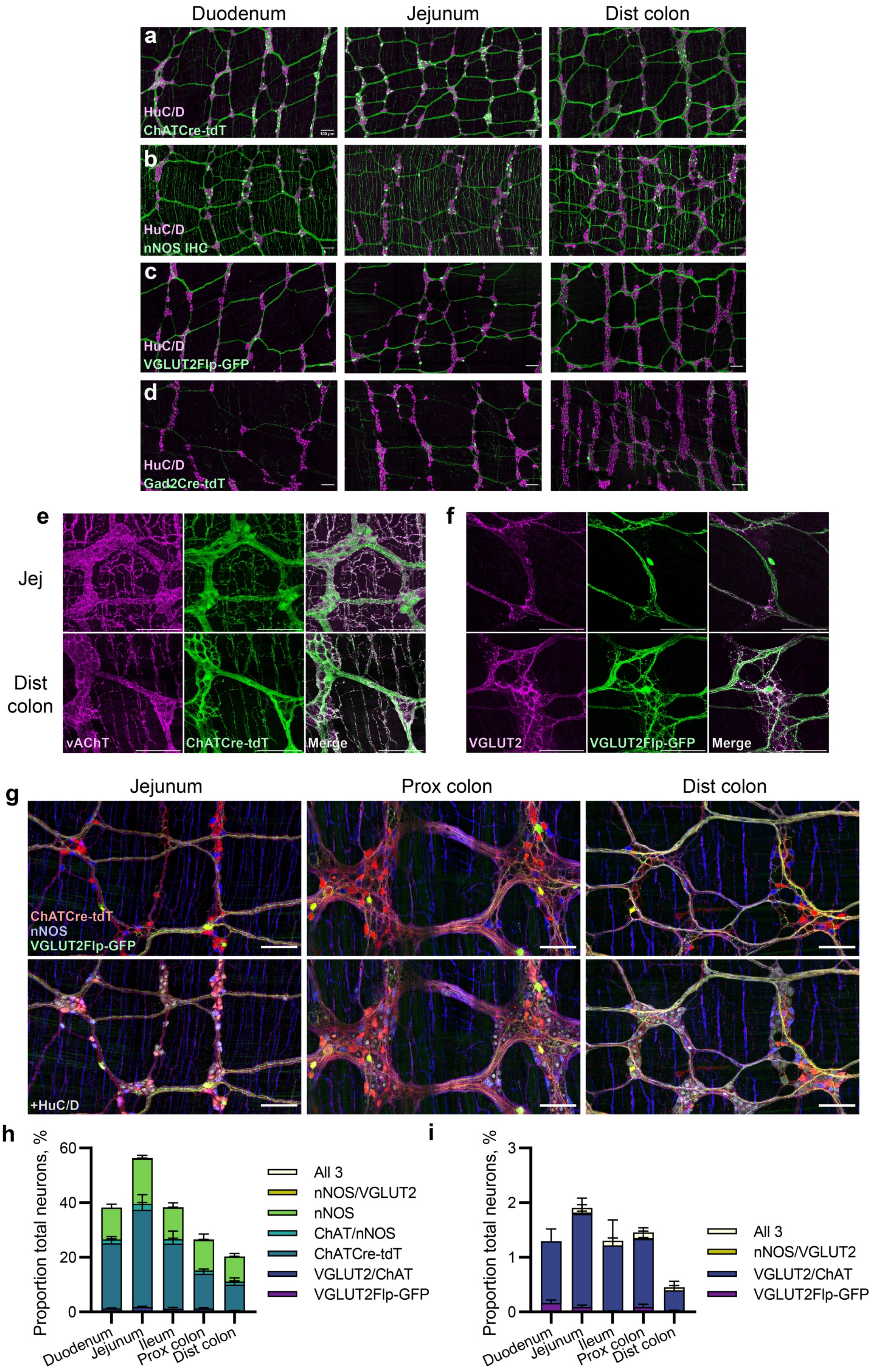

Duodenum
Jejunum
Dist colon
a
HuC/D
ChATCre-tdT
b
HuC/D
nNOS IHC
c
HuC/D
VGLUT2Flp-GFP
d
HuC/D
Gad2Cre-tdT
e
Jej
Dist colon
vAChT
ChATCre-tdT
Merge
f
VGLUT2
VGLUT2Flp-GFP
Merge
g
Jejunum
Prox colon
Dist colon
ChATCre-tdT
nNOS
VGLUT2Flp-GFP
+HuC/D
h
Proportion total neurons, %
60
40
20
0
Duodenum
Jejunum
Ileum
Prox colon
Dist colon
All 3
nNOS/VGLUT2
nNOS
ChAT/nNOS
ChATCre-tdT
VGLUT2/ChAT
VGLUT2Flp-GFP
i
Proportion total neurons, %
3
2
1
0
Duodenum
Jejunum
Ileum
Prox colon
Dist colon
All 3
nNOS/VGLUT2
VGLUT2/ChAT
VGLUT2Flp-GFP

**Supplementary Fig. 3 (Supporting Figure 3) | Neurotransmitter distribution in the ENS. a-d,** Immunohistochemical labelling of adult wholemount MP duodenum, jejunum and DC for HuC/D (magenta) and ChATCre-tdT (**a**), nNOS (**b**), VGLUT2Flp-GFP (**c**) or Gad2Cre-tdT (**d**) (green). **e**, Immunohistochemical labelling of adult ChATCre-tdT (green) wholemount MP jejunum (top) and DC (bottom) for cholinergic marker vAChT (magenta). **f**, Immunohistochemical staining of adult VGLUT2Flp-GFP (green) wholemount jejunum (top) and DC (bottom) MP for VGLUT2 (magenta). **g**, Immunohistochemical labelling of adult VGLUT2Flp-GFP(green)-ChATCre-tdT (red) wholemount MP jejunum, PC and DC for nNOS (blue), with (bottom) or without (top) HuC/D (gray), allowing concomitant visualization of three major enteric neurotransmitters. **h**, Proportion of total HuC/D neurons (mean ± SEM) across intestinal regions positive for VGLUT2Flp-GFP, ChATCre-tdT and nNOS, alone or in combination with each other. n = 5 mice. **i,** Same data as (**h**) but with the largest three groups removed (nNOS alone, ChATCre-tdT alone, and nNOS/ChATCre-tdT) to allow easier visualization of less frequent groups. Scale bars 100 µm. ChAT, choline acetyltransferase; Cre, Cre recombinase; DC, distal colon; Flp, Flp recombinase; Gad2, glutamate decarboxylase 2; GFP, green fluorescent protein; IHC, immunohistochemistry; MP, myenteric plexus; nNOS, neuronal nitric oxide synthase; PC, proximal colon; tdT, tdTomato; vAChT, vesicular acetylcholine transporter; VGLUT2, vesicular glutamate transporter 2.
