## Supplementary Fig. 4 for "Regional cytoarchitecture of the adult and developing mouse enteric nervous system"

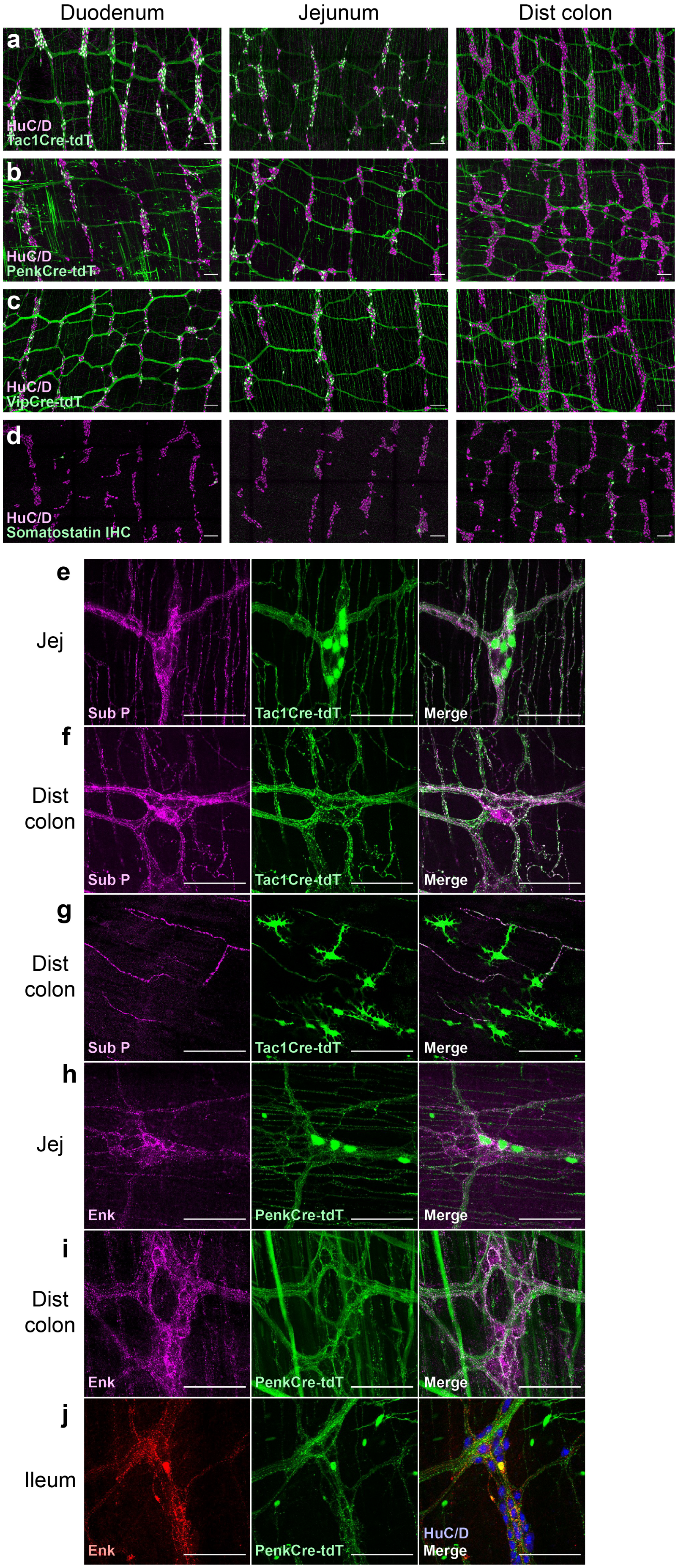

Duodenum
Jejunum
Dist colon
a
HuC/D
Tac1Cre-tdT
b
HuC/D
PenkCre-tdT
c
HuC/D
VipCre-tdT
d
HuC/D
Somatostatin IHC
e
Jej
Sub P
Tac1Cre-tdT
Merge
f
Dist colon
Sub P
Tac1Cre-tdT
Merge
g
Dist colon
Sub P
Tac1Cre-tdT
Merge
h
Jej
Enk
PenkCre-tdT
Merge
i
Dist colon
Enk
PenkCre-tdT
Merge
j
Ileum
Enk
PenkCre-tdT
HuC/D
Merge

**Supplementary Fig. 4 (Supporting Figure 3) | Neuropeptide distribution in the ENS. a-d**, Immunohistochemical labelling of adult wholemount MP duodenum, jejunum and DC for HuC/D (magenta) and Tac1Cre-tdT (a), PenkCre-tdT (b), VipCre-tdT (c) or somatostatin (d) (green). e,f, Immunohistochemical labelling of adult Tac1Cre-tdT (green) wholemount MP jejunum (e) and DC (f) for substance P (magenta). g, Representative images of non-neuronal cells positive for the Tac1Cre-tdT reporter (green), alongside substance P IHC (magenta). h,i, Immunohistochemical labelling of adult PenkCre-tdT (green) wholemount MP jejunum (e) and DC (f) for enkephalin (magenta). j, Representative images of a non-neuronal cell positive for both the PenkCre-tdT reporter (green) and enkephalin IHC (red), but negative for HuC/D (blue). Scale bars 100 µm. Cre, Cre recombinase; DC, distal colon; Enk, enkephalin; IHC, immunohistochemistry; MP, myenteric plexus; Penk, proenkephalin; Sub P, substance P; Tac1, Tachykinin Precursor 1; tdT, tdTomato; Vip, vasoactive intestinal peptide.
