## Supplementary Fig. 5 for "Regional cytoarchitecture of the adult and developing mouse enteric nervous system"

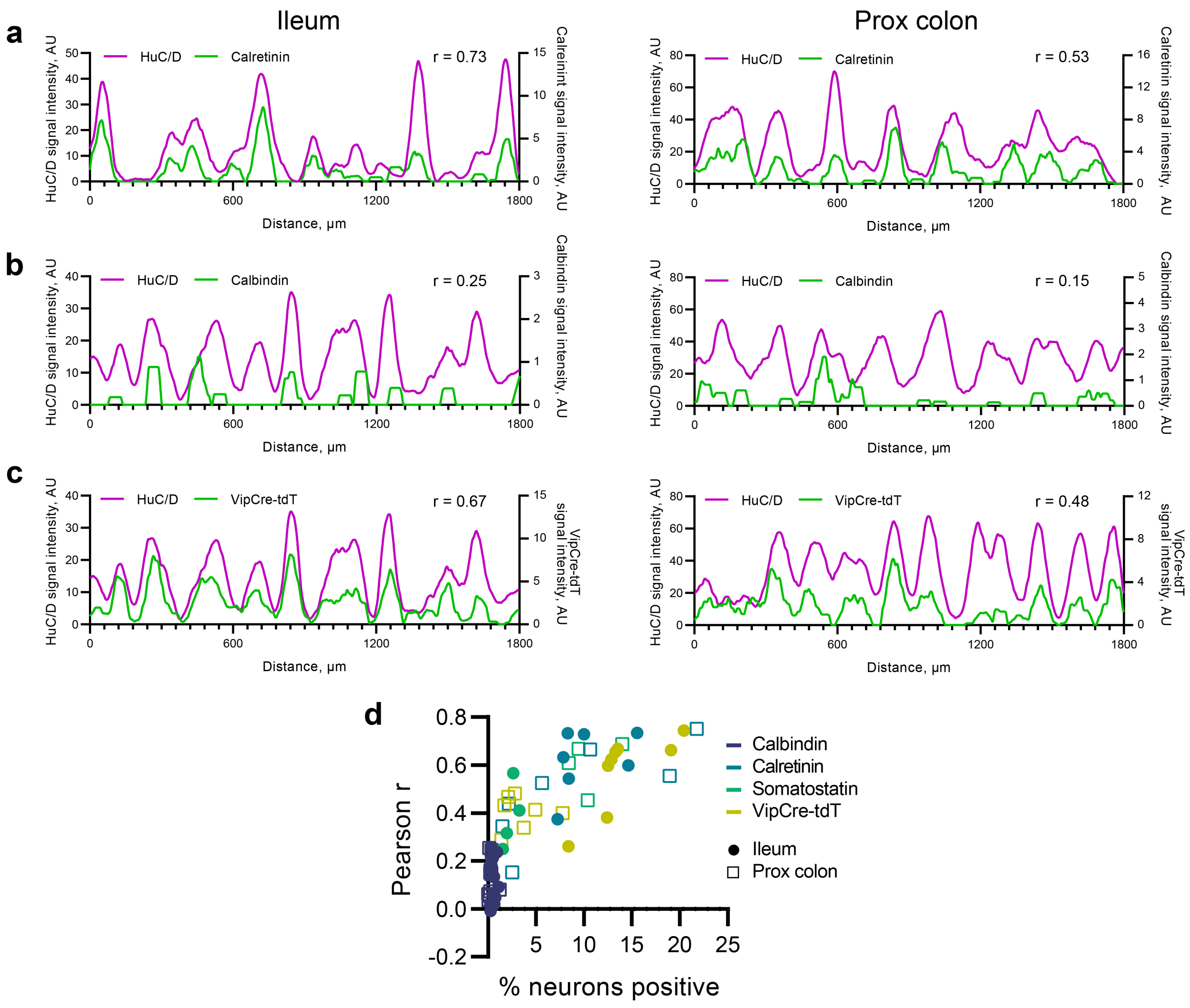


**Supplementary Fig. 5 (Supporting Figure 3) | Neuronal subtypes are evenly distributed across enteric neuronal stripes. a-c**, Representative smoothed profiles of HuC/D and calretinin (**a**), calbindin (**b**), or VipCre-tdT (**c**) signal intensity along the longitudinal axis in ileum (left) and PC (right). r value indicates Pearson r correlation between the two profiles. **d**, Scatter plot showing the relationship between the proportion of total (HuC/D) neurons positive for a given marker (calbindin, calretinin, somatostatin or VipCre-tdT) in the ileum or PC and the Pearson r value for that sample, representing the strength of correlation between the HuC/D signal intensity profile and that of the marker (as in **a-c**). Note that the association is very weak for lowly expressed markers (e.g. calbindin) but rises steeply when more than 2-3% of neurons are positive for a given marker. Cre, Cre recombinase; PC, proximal colon; tdT, tdTomato; Vip, vasoactive intestinal peptide.
