## Supplementary Table 1 for "Regional cytoarchitecture of the adult and developing mouse enteric nervous system"

**Fig 1h Stripe width**

**Repeated measures ANOVA summary**

| | |
|---|---|
| Assume sphericity? | No |
| F | 36.17 |
| P value | <0.0001 |
| P value summary | **** |
| Statistically significant (P < 0.05)? | Yes |

| **ANOVA table** | **SS** | **DF** | **MS** | **F (DFn, DFd)** | **P value** |
|---|---|---|---|---|---|
| Treatment (between columns) | 7122 | 4 | 1781 | F (3.436, 113.4) = 36.17 | P<0.0001 |
| Individual (between rows) | 1804 | 33 | 54.66 | F (33, 132) = 1.110 | P=0.3307 |
| Residual (random) | 6498 | 132 | 49.23 | | |
| Total | 15424 | 169 | | | |

| **Tukey's multiple comparisons test** | **Mean Diff.** | **95.00% CI of diff.** | **Summary** | **Adjusted P Value** |
|---|---|---|---|---|
| D vs. J | -3.678 | -8.532 to 1.176 | ns | 0.2102 |
| D vs. I | -9.736 | -14.41 to -5.058 | **** | <0.0001 |
| D vs. PC | -18.9 | -24.22 to -13.57 | **** | <0.0001 |
| D vs. DC | -5.553 | -9.716 to -1.391 | ** | 0.0044 |
| J vs. I | -6.058 | -11.35 to -0.7648 | * | 0.0184 |
| J vs. PC | -15.22 | -19.67 to -10.76 | **** | <0.0001 |
| J vs. DC | -1.875 | -5.700 to 1.950 | ns | 0.6232 |
| I vs. PC | -9.159 | -14.76 to -3.555 | *** | 0.0004 |
| I vs. DC | 4.183 | -1.408 to 9.774 | ns | 0.2207 |
| PC vs. DC | 13.34 | 8.380 to 18.30 | **** | <0.0001 |

**Fig 1h Iinterstripe distance**

| | | | | |
|---|---|---|---|---|
| **Mixed-effects model (REML)** | Matching: Across row | | | |
| Assume sphericity? | No | | | |
| Alpha | 0.05 | | | |
| **Fixed effect (type III)** | **P value** | **P value summary** | **Statistically signific** | **F (DFn, DFd)** |
| Treatment (between columns) | 0.0002 | *** | Yes | F (3.231, 97.73) = 6.915 |

| **Tukey's multiple comparisons test** | **Mean Diff.** | **95.00% CI of diff.** | **Summary** | **Adjusted P Value** |
|---|---|---|---|---|
| D vs. J | -9.881 | -68.21 to 48.45 | ns | 0.9857 |
| D vs. I | -6.594 | -55.70 to 42.51 | ns | 0.9944 |
| D vs. PC | 45.04 | 7.388 to 82.70 | * | 0.0138 |
| D vs. DC | 53.81 | -3.206 to 110.8 | ns | 0.0707 |
| J vs. I | 3.287 | -60.58 to 67.16 | ns | 0.9999 |
| J vs. PC | 54.92 | 6.801 to 103.0 | * | 0.0205 |
| J vs. DC | 63.69 | -0.5408 to 127.9 | ns | 0.0526 |
| I vs. PC | 51.64 | 14.61 to 88.66 | ** | 0.0032 |
| I vs. DC | 60.41 | 11.51 to 109.3 | * | 0.0102 |
| PC vs. DC | 8.769 | -24.94 to 42.47 | ns | 0.9385 |

Fig 1j MP density

**Repeated measures ANOVA summary**

| | |
|---|---|
| Assume sphericity? | No |
| F | 130.9 |
| P value | <0.0001 |
| P value summary | **** |
| Statistically significant (P < 0.05)? | Yes |

| **ANOVA table** | **SS** | **DF** | **MS** | **F (DFn, DFd)** | **P value** |
|---|---|---|---|---|---|
| Treatment (between columns) | 3393734 | 4 | 848433 | F (2.155, 64.65) = 130.9 | P<0.0001 |
| Individual (between rows) | 304684 | 30 | 10156 | F (30, 120) = 1.567 | P=0.0469 |
| Residual (random) | 777665 | 120 | 6481 | | |
| Total | 4476083 | 154 | | | |

| **Tukey's multiple comparisons test** | **Mean Diff.** | **95.00% CI of diff.** | **Summary** | **Adjusted P Value** |
|---|---|---|---|---|
| Duodenum vs. Jejunum | -29.09 | -58.58 to 0.3929 | ns | 0.0545 |
| Duodenum vs. Ileum | -115.5 | -148.6 to -82.36 | **** | <0.0001 |
| Duodenum vs. Prox colon | -413 | -484.5 to -341.5 | **** | <0.0001 |
| Duodenum vs. Dist colon | -194.8 | -242.9 to -146.8 | **** | <0.0001 |
| Jejunum vs. Ileum | -86.39 | -117.1 to -55.72 | **** | <0.0001 |
| Jejunum vs. Prox colon | -383.9 | -461.8 to -306.0 | **** | <0.0001 |
| Jejunum vs. Dist colon | -165.7 | -214.7 to -116.8 | **** | <0.0001 |
| Ileum vs. Prox colon | -297.5 | -373.9 to -221.1 | **** | <0.0001 |
| Ileum vs. Dist colon | -79.36 | -135.1 to -23.61 | ** | 0.0023 |
| Prox colon vs. Dist colon | 218.1 | 131.9 to 304.4 | **** | <0.0001 |

Fig 1k SMP density

**Repeated measures ANOVA summary**

| | |
|---|---|
| Assume sphericity? | No |
| F | 15.92 |
| P value | 0.001 |
| P value summary | ** |
| Statistically significant (P < 0.05)? | Yes |

| **ANOVA table** | **SS** | **DF** | **MS** | **F (DFn, DFd)** | **P value** |
|---|---|---|---|---|---|
| Treatment (between columns) | 28079 | 4 | 7020 | F (1.680, 10.08) = 15.92 | P=0.0010 |
| Individual (between rows) | 2219 | 6 | 369.9 | F (6, 24) = 0.8389 | P=0.5523 |
| Residual (random) | 10582 | 24 | 440.9 | | |
| Total | 40881 | 34 | | | |

| **Tukey's multiple comparisons test** | **Mean Diff.** | **95.00% CI of diff.** | **Summary** | **Adjusted P Value** |
|---|---|---|---|---|
| Duodenum vs. Jejunum | 36.22 | 0.8690 to 71.58 | * | 0.0452 |
| Duodenum vs. Ileum | 61.5 | 20.38 to 102.6 | ** | 0.0078 |
| Duodenum vs. Prox colon | 40.19 | -26.58 to 107.0 | ns | 0.275 |
| Duodenum vs. Dist colon | 85.18 | 28.20 to 142.2 | ** | 0.0079 |
| Jejunum vs. Ileum | 25.28 | 9.024 to 41.53 | ** | 0.0064 |
| Jejunum vs. Prox colon | 3.968 | -42.25 to 50.18 | ns | 0.997 |
| Jejunum vs. Dist colon | 48.96 | 10.42 to 87.50 | * | 0.0174 |
| Ileum vs. Prox colon | -21.31 | -61.64 to 19.02 | ns | 0.3725 |
| Ileum vs. Dist colon | 23.68 | -5.937 to 53.31 | ns | 0.1171 |
| Prox colon vs. Dist colon | 44.99 | 18.69 to 71.29 | ** | 0.0039 |

**Fig 2e SI length**

| **Ordinary one-way ANOVA summary** | |
|---|---|
| F | 1733 |
| P value | <0.0001 |
| P value summary | **** |
| Significant diff. among means (P < 0.05)? | Yes |
| R squared | 0.9939 |

| **ANOVA table** | **SS** | **DF** | **MS** | **F (DFn, DFd)** | **P value** |
|---|---|---|---|---|---|
| Treatment (between columns) | 1366 | 5 | 273.2 | F (5, 53) = 1733 | P<0.0001 |
| Residual (within columns) | 8.356 | 53 | 0.1577 | | |
| Total | 1375 | 58 | | | |

| **Dunnett's multiple comparisons test** | **Mean Diff.** | **95.00% CI of diff.** | **Summary** | **Adjusted P Value** |
|---|---|---|---|---|
| P21 vs. E14.5 | 18.11 | 17.49 to 18.72 | **** | <0.0001 |
| P21 vs. E16.5 | 15.59 | 14.98 to 16.21 | **** | <0.0001 |
| P21 vs. E18.5 | 13.93 | 13.30 to 14.56 | **** | <0.0001 |
| P21 vs. P0 | 13.25 | 12.59 to 13.92 | **** | <0.0001 |
| P21 vs. P10 | 4.983 | 4.292 to 5.674 | **** | <0.0001 |

**Fig 2f colon length**

| Ordinary one-way ANOVA summary | |
|---|---|
| F | 233.4 |
| P value | <0.0001 |
| P value summary | **** |
| Significant diff. among means (P < 0.05)? | Yes |
| R squared | 0.9565 |

| ANOVA table | SS | DF | MS | F (DFn, DFd) | P value |
|---|---|---|---|---|---|
| Treatment (between columns) | 47.92 | 5 | 9.584 | F (5, 53) = 233.4 | P<0.0001 |
| Residual (within columns) | 2.177 | 53 | 0.04107 | | |
| Total | 50.1 | 58 | | | |

| Dunnett's multiple comparisons test | Mean Diff. | 95.00% CI of diff. | Summary | Adjusted P Value |
|---|---|---|---|---|
| P21 vs. E14.5 | 3.287 | 2.971 to 3.602 | **** | <0.0001 |
| P21 vs. E16.5 | 2.953 | 2.638 to 3.269 | **** | <0.0001 |
| P21 vs. E18.5 | 2.6 | 2.278 to 2.922 | **** | <0.0001 |
| P21 vs. P0 | 2.413 | 2.075 to 2.750 | **** | <0.0001 |
| P21 vs. P10 | 0.8333 | 0.4806 to 1.186 | **** | <0.0001 |

**Fig 2g jejunum density**

**Ordinary one-way ANOVA summary**

| | |
|---|---|
| F | 19.94 |
| P value | <0.0001 |
| P value summary | **** |
| Significant diff. among means (P < 0.05)? | Yes |
| R squared | 0.7401 |

| ANOVA table | SS | DF | MS | F (DFn, DFd) | P value |
|---|---|---|---|---|---|
| Treatment (between columns) | 7971403 | 5 | 1594281 | F (5, 35) = 19.94 | P<0.0001 |
| Residual (within columns) | 2798676 | 35 | 79962 | | |
| Total | 10770079 | 40 | | | |

| Dunnett's multiple comparisons test | Mean Diff. | 95.00% CI of diff. | Summary | Adjusted P Value |
|---|---|---|---|---|
| P21 vs. E14.5 | -1354 | -1833 to -874.0 | **** | <0.0001 |
| P21 vs. E16.5 | -1279 | -1747 to -810.1 | **** | <0.0001 |
| P21 vs. E18.5 | -929.7 | -1455 to -404.3 | *** | 0.0003 |
| P21 vs. P0 | -1073 | -1570 to -576.7 | **** | <0.0001 |
| P21 vs. P10 | -299.4 | -808.0 to 209.3 | ns | 0.3582 |

**Fig 2h DC density**

| Ordinary one-way ANOVA summary | |
|---|---|
| F | 19.65 |
| P value | <0.0001 |
| P value summary | **** |
| Significant diff. among means (P < 0.05)? | Yes |
| R squared | 0.7844 |

| ANOVA table | SS | DF | MS | F (DFn, DFd) | P value |
|---|---|---|---|---|---|
| Treatment (between columns) | 21201106 | 5 | 4240221 | F (5, 27) = 19.65 | P<0.0001 |
| Residual (within columns) | 5825854 | 27 | 215772 | | |
| Total | 27026960 | 32 | | | |

| Dunnett's multiple comparisons test | Mean Diff. | 95.00% CI of diff. | Summary | Adjusted P Value |
|---|---|---|---|---|
| P21 vs. E14.5 | 335.8 | -603.7 to 1275 | ns | 0.7843 |
| P21 vs. E16.5 | -1938 | -2665 to -1210 | **** | <0.0001 |
| P21 vs. E18.5 | -1660 | -2530 to -790.7 | *** | 0.0001 |
| P21 vs. P0 | -1591 | -2385 to -797.3 | **** | <0.0001 |
| P21 vs. P10 | -759.8 | -1554 to 34.23 | ns | 0.0639 |

**Fig 2i jejunum interstripe**

| **Ordinary one-way ANOVA summary** | |
|---|---|
| F | 9.202 |
| P value | 0.0008 |
| P value summary | *** |
| Significant diff. among means (P < 0.05)? | Yes |
| R squared | 0.6189 |

| **ANOVA table** | **SS** | **DF** | **MS** | **F (DFn, DFd)** | **P value** |
|---|---|---|---|---|---|
| Treatment (between columns) | 23003 | 3 | 7668 | F (3, 17) = 9.202 | P=0.0008 |
| Residual (within columns) | 14166 | 17 | 833.3 | | |
| Total | 37168 | 20 | | | |

| **Dunnett's multiple comparisons test** | **Mean Diff.** | **95.00% CI of diff.** | **Summary** | **Adjusted P Value** |
|---|---|---|---|---|
| P21 vs. E18.5 | 81.24 | 28.02 to 134.5 | ** | 0.0032 |
| P21 vs. P0 | 59.66 | 9.364 to 109.9 | * | 0.0195 |
| P21 vs. P10 | 5.817 | -45.72 to 57.35 | ns | 0.9783 |

**Fig 2j DC interstripe**

| **Ordinary one-way ANOVA summary** | |
|---|---|
| F | 34.51 |
| P value | <0.0001 |
| P value summary | **** |
| Significant diff. among means (P < 0.05)? | Yes |
| R squared | 0.8625 |

| **ANOVA table** | **SS** | **DF** | **MS** | **F (DFn, DFd)** | **P value** |
|---|---|---|---|---|---|
| Treatment (between columns) | 60588 | 2 | 30294 | F (2, 11) = 34.51 | P<0.0001 |
| Residual (within columns) | 9657 | 11 | 877.9 | | |
| Total | 70245 | 13 | | | |

| **Dunnett's multiple comparisons test** | **Mean Diff.** | **95.00% CI of diff.** | **Summary** | **Adjusted P Value** |
|---|---|---|---|---|
| P21 vs. P0 | 161.1 | 111.0 to 211.2 | **** | <0.0001 |
| P21 vs. P10 | 59.46 | 9.334 to 109.6 | * | 0.022 |

**Fig 3 Calbindin**

**Repeated measures ANOVA summary**

| | |
|---|---|
| Assume sphericity? | No |
| F | 7.284 |
| P value | 0.001 |
| P value summary | *** |
| Statistically significant (P < 0.05)? | Yes |

| **ANOVA table** | **SS** | **DF** | **MS** | **F (DFn, DFd)** | **P value** |
|---|---|---|---|---|---|
| Treatment (between columns) | 2.509 | 4 | 0.6272 | F (2.647, 34.42) = 7.284 | P=0.0010 |
| Individual (between rows) | 1.773 | 13 | 0.1364 | F (13, 52) = 1.584 | P=0.1203 |
| Residual (random) | 4.477 | 52 | 0.0861 | | |
| Total | 8.759 | 69 | | | |

| **Tukey's multiple comparisons test** | **Mean Diff.** | **95.00% CI of diff.** | **Summary** | **Adjusted P Value** |
|---|---|---|---|---|
| D vs. J | -0.1387 | -0.5297 to 0.2523 | ns | 0.7951 |
| D vs. I | -0.05037 | -0.4680 to 0.3673 | ns | 0.995 |
| D vs. PC | 0.2181 | -0.01846 to 0.4546 | ns | 0.0767 |
| D vs. DC | 0.3782 | -0.02369 to 0.7801 | ns | 0.0691 |
| J vs. I | 0.08832 | -0.2194 to 0.3960 | ns | 0.8904 |
| J vs. PC | 0.3568 | -0.04480 to 0.7583 | ns | 0.0918 |
| J vs. DC | 0.5169 | 0.2338 to 0.7999 | *** | 0.0005 |
| I vs. PC | 0.2685 | -0.1122 to 0.6491 | ns | 0.2319 |
| I vs. DC | 0.4286 | 0.1545 to 0.7026 | ** | 0.0021 |
| PC vs. DC | 0.1601 | -0.1844 to 0.5046 | ns | 0.6013 |

**Fig 3 Calretinin**

**Repeated measures ANOVA summary**

| | |
|---|---|
| Assume sphericity? | No |
| F | 10.64 |
| P value | 0.0003 |
| P value summary | *** |
| Statistically significant (P < 0.05)? | Yes |

| ANOVA table | SS | DF | MS | F (DFn, DFd) | P value |
|---|---|---|---|---|---|
| Treatment (between columns) | 1562 | 4 | 390.6 | F (2.542, 20.34) = 10.64 | P=0.0003 |
| Individual (between rows) | 160.9 | 8 | 20.11 | F (8, 32) = 0.5479 | P=0.8114 |
| Residual (random) | 1174 | 32 | 36.7 | | |
| Total | 2898 | 44 | | | |

| Tukey's multiple comparisons test | Mean Diff. | 95.00% CI of diff. | Summary | Adjusted P Value |
|---|---|---|---|---|
| D vs. J | -9.355 | -19.33 to 0.6201 | ns | 0.067 |
| D vs. I | 2.416 | -5.052 to 9.885 | ns | 0.7938 |
| D vs. PC | 4.03 | -5.316 to 13.38 | ns | 0.595 |
| D vs. DC | 8.362 | -1.962 to 18.69 | ns | 0.1226 |
| J vs. I | 11.77 | 2.180 to 21.36 | * | 0.0176 |
| J vs. PC | 13.39 | -0.6182 to 27.39 | ns | 0.0616 |
| J vs. DC | 17.72 | 8.486 to 26.95 | ** | 0.0011 |
| I vs. PC | 1.614 | -8.230 to 11.46 | ns | 0.9765 |
| I vs. DC | 5.946 | 1.495 to 10.40 | * | 0.0109 |
| PC vs. DC | 4.332 | -7.250 to 15.91 | ns | 0.7031 |

**Fig 3 Secretagogin**

**Repeated measures ANOVA summary**

| | |
|---|---|
| Assume sphericity? | No |
| F | 9.885 |
| P value | 0.0001 |
| P value summary | *** |
| Statistically significant (P < 0.05)? | Yes |

| **ANOVA table** | **SS** | **DF** | **MS** | **F (DFn, DFd)** | **P value** |
|---|---|---|---|---|---|
| Treatment (between columns) | 97.67 | 4 | 24.42 | F (2.681, 34.85) = 9.885 | P=0.0001 |
| Individual (between rows) | 71.63 | 13 | 5.51 | F (13, 52) = 2.231 | P=0.0208 |
| Residual (random) | 128.5 | 52 | 2.47 | | |
| Total | 297.8 | 69 | | | |

| **Tukey's multiple comparisons test** | **Mean Diff.** | **95.00% CI of diff.** | **Summary** | **Adjusted P Value** |
|---|---|---|---|---|
| D vs. J | -0.8372 | -2.629 to 0.9544 | ns | 0.5966 |
| D vs. I | -1.45 | -4.021 to 1.122 | ns | 0.4267 |
| D vs. PC | 1.588 | 0.1902 to 2.986 | * | 0.0233 |
| D vs. DC | 1.314 | -0.3060 to 2.935 | ns | 0.1377 |
| J vs. I | -0.6123 | -2.393 to 1.168 | ns | 0.8121 |
| J vs. PC | 2.426 | 0.5323 to 4.319 | * | 0.0103 |
| J vs. DC | 2.152 | 0.6824 to 3.621 | ** | 0.0037 |
| I vs. PC | 3.038 | 0.9463 to 5.129 | ** | 0.0039 |
| I vs. DC | 2.764 | 0.5140 to 5.014 | * | 0.0138 |
| PC vs. DC | -0.2739 | -1.768 to 1.220 | ns | 0.9762 |

**Fig 3 ChATCre-tdT**

**Repeated measures ANOVA summary**

| | |
|---|---|
| Assume sphericity? | No |
| F | 15.34 |
| P value | 0.0001 |
| P value summary | *** |
| Statistically significant (P < 0.05)? | Yes |

| **ANOVA table** | **SS** | **DF** | **MS** | **F (DFn, DFd)** | **P value** |
|---|---|---|---|---|---|
| Treatment (between columns) | 5915 | 4 | 1479 | F (2.185, 17.48) = 15.34 | P=0.0001 |
| Individual (between rows) | 1176 | 8 | 147 | F (8, 32) = 1.525 | P=0.1876 |
| Residual (random) | 3084 | 32 | 96.38 | | |
| Total | 10176 | 44 | | | |

| **Tukey's multiple comparisons test** | **Mean Diff.** | **95.00% CI of diff.** | **Summary** | **Adjusted P Value** |
|---|---|---|---|---|
| D vs. J | -14.47 | -25.60 to -3.333 | * | 0.0128 |
| D vs. I | 3.546 | -14.96 to 22.05 | ns | 0.9593 |
| D vs. PC | 17.14 | 4.501 to 29.78 | ** | 0.01 |
| D vs. DC | 15.31 | 7.454 to 23.17 | ** | 0.001 |
| J vs. I | 18.01 | -5.276 to 41.30 | ns | 0.1454 |
| J vs. PC | 31.6 | 15.13 to 48.08 | ** | 0.0011 |
| J vs. DC | 29.78 | 18.57 to 40.99 | *** | 0.0001 |
| I vs. PC | 13.59 | -2.939 to 30.12 | ns | 0.1157 |
| I vs. DC | 11.76 | -9.104 to 32.63 | ns | 0.367 |
| PC vs. DC | -1.829 | -16.67 to 13.01 | ns | 0.9918 |

**Fig 3 nNOS**

**Repeated measures ANOVA summary**

| | |
|---|---|
| Assume sphericity? | No |
| F | 5.236 |
| P value | 0.006 |
| P value summary | ** |
| Statistically significant (P < 0.05)? | Yes |

| ANOVA table | SS | DF | MS | F (DFn, DFd) | P value |
|---|---|---|---|---|---|
| Treatment (between columns) | 456.6 | 4 | 114.1 | F (2.832, 28.32) = 5.236 | P=0.0060 |
| Individual (between rows) | 445 | 10 | 44.5 | F (10, 40) = 2.041 | P=0.0541 |
| Residual (random) | 872 | 40 | 21.8 | | |
| Total | 1774 | 54 | | | |

| Tukey's multiple comparisons test | Mean Diff. | 95.00% CI of diff. | Summary | Adjusted P Value |
|---|---|---|---|---|
| D vs. J | -5.154 | -9.096 to -1.212 | * | 0.0106 |
| D vs. I | 1.088 | -5.705 to 7.882 | ns | 0.9824 |
| D vs. PC | 2.114 | -4.224 to 8.452 | ns | 0.804 |
| D vs. DC | 3.09 | -3.046 to 9.226 | ns | 0.4977 |
| J vs. I | 6.242 | 0.05728 to 12.43 | * | 0.0477 |
| J vs. PC | 7.268 | 0.9277 to 13.61 | * | 0.0237 |
| J vs. DC | 8.244 | 2.440 to 14.05 | ** | 0.0061 |
| I vs. PC | 1.026 | -7.273 to 9.325 | ns | 0.9933 |
| I vs. DC | 2.002 | -6.744 to 10.75 | ns | 0.9383 |
| PC vs. DC | 0.9759 | -4.707 to 6.659 | ns | 0.9773 |

**Fig 3 VGLUT2Flp-GFP**

| Repeated measures ANOVA summary | |
|---|---|
| Assume sphericity? | No |
| F | 11.47 |
| P value | <0.0001 |
| P value summary | **** |
| Statistically significant (P < 0.05)? | Yes |

| ANOVA table | SS | DF | MS | F (DFn, DFd) | P value |
|---|---|---|---|---|---|
| Treatment (between columns) | 23.94 | 4 | 5.986 | F (2.325, 58.12) = 11.47 | P<0.0001 |
| Individual (between rows) | 17.74 | 25 | 0.7095 | F (25, 100) = 1.359 | P=0.1450 |
| Residual (random) | 52.2 | 100 | 0.522 | | |
| Total | 93.88 | 129 | | | |

| Tukey's multiple comparisons test | Mean Diff. | 95.00% CI of diff. | Summary | Adjusted P Value |
|---|---|---|---|---|
| D vs. J | -0.3991 | -0.8439 to 0.04559 | ns | 0.0939 |
| D vs. I | -0.02846 | -0.4867 to 0.4298 | ns | 0.9997 |
| D vs. PC | -0.2395 | -0.8966 to 0.4176 | ns | 0.8197 |
| D vs. DC | 0.8424 | 0.4942 to 1.191 | **** | <0.0001 |
| J vs. I | 0.3707 | -0.1774 to 0.9188 | ns | 0.3014 |
| J vs. PC | 0.1596 | -0.6484 to 0.9676 | ns | 0.9768 |
| J vs. DC | 1.242 | 0.8651 to 1.618 | **** | <0.0001 |
| I vs. PC | -0.211 | -1.005 to 0.5831 | ns | 0.9339 |
| I vs. DC | 0.8708 | 0.4445 to 1.297 | **** | <0.0001 |
| PC vs. DC | 1.082 | 0.3104 to 1.853 | ** | 0.0031 |

**Fig 3 Gad2Cre-tdT**

**Repeated measures ANOVA summary**

| | |
|---|---|
| Assume sphericity? | No |
| F | 26.29 |
| P value | 0.0015 |
| P value summary | ** |
| Statistically significant (P < 0.05)? | Yes |

| **ANOVA table** | **SS** | **DF** | **MS** | **F (DFn, DFd)** | **P value** |
|---|---|---|---|---|---|
| Treatment (between columns) | 2.231 | 4 | 0.5578 | F (1.873, 5.618) = 26.29 | P=0.0015 |
| Individual (between rows) | 0.1174 | 3 | 0.03913 | F (3, 12) = 1.845 | P=0.1928 |
| Residual (random) | 0.2546 | 12 | 0.02121 | | |
| Total | 2.603 | 19 | | | |

| **Tukey's multiple comparisons test** | **Mean Diff.** | **95.00% CI of diff.** | **Summary** | **Adjusted P Value** |
|---|---|---|---|---|
| D vs. J | -0.04703 | -0.4853 to 0.3912 | ns | 0.9714 |
| D vs. I | -0.754 | -1.338 to -0.1703 | * | 0.0248 |
| D vs. PC | 0.117 | 0.06837 to 0.1657 | ** | 0.0041 |
| D vs. DC | 0.1649 | -0.3543 to 0.6841 | ns | 0.545 |
| J vs. I | -0.707 | -1.354 to -0.06012 | * | 0.0393 |
| J vs. PC | 0.164 | -0.3011 to 0.6292 | ns | 0.4737 |
| J vs. DC | 0.2119 | -0.1226 to 0.5464 | ns | 0.1568 |
| I vs. PC | 0.8711 | 0.2486 to 1.494 | * | 0.0198 |
| I vs. DC | 0.9189 | 0.04956 to 1.788 | * | 0.0431 |
| PC vs. DC | 0.04785 | -0.4906 to 0.5863 | ns | 0.9852 |

**Fig 3 Tac1Cre-tdT**

**Repeated measures ANOVA summary**

| | |
|---|---|
| Assume sphericity? | No |
| F | 17.56 |
| P value | 0.0004 |
| P value summary | *** |
| Statistically significant (P < 0.05)? | Yes |

| **ANOVA table** | **SS** | **DF** | **MS** | **F (DFn, DFd)** | **P value** |
|---|---|---|---|---|---|
| Treatment (between columns) | 6568 | 4 | 1642 | F (2.089, 10.45) = 17.56 | P=0.0004 |
| Individual (between rows) | 813.3 | 5 | 162.7 | F (5, 20) = 1.740 | P=0.1716 |
| Residual (random) | 1870 | 20 | 93.5 | | |
| Total | 9251 | 29 | | | |

| **Tukey's multiple comparisons test** | **Mean Diff.** | **95.00% CI of diff.** | **Summary** | **Adjusted P Value** |
|---|---|---|---|---|
| D vs. J | 4.278 | -15.28 to 23.83 | * | 0.0106 |
| D vs. I | 8.392 | -18.87 to 35.65 | ns | 0.9824 |
| D vs. PC | 32.14 | 16.72 to 47.57 | ns | 0.804 |
| D vs. DC | 35.57 | 17.21 to 53.92 | ns | 0.4977 |
| J vs. I | 4.114 | -21.70 to 29.93 | * | 0.0477 |
| J vs. PC | 27.87 | 9.294 to 46.44 | * | 0.0237 |
| J vs. DC | 31.29 | 13.49 to 49.08 | ** | 0.0061 |
| I vs. PC | 23.75 | -8.634 to 56.14 | ns | 0.9933 |
| I vs. DC | 27.17 | -2.679 to 57.03 | ns | 0.9383 |
| PC vs. DC | 3.421 | -3.424 to 10.27 | ns | 0.9773 |

**Fig 3 PenkCre-tdT**

**Repeated measures ANOVA summary**

| | |
|---|---|
| Assume sphericity? | No |
| F | 18.77 |
| P value | <0.0001 |
| P value summary | **** |
| Statistically significant (P < 0.05)? | Yes |

| **ANOVA table** | **SS** | **DF** | **MS** | **F (DFn, DFd)** | **P value** |
|---|---|---|---|---|---|
| Treatment (between columns) | 1016 | 4 | 253.9 | F (2.245, 20.21) = 18.77 | P<0.0001 |
| Individual (between rows) | 44.22 | 9 | 4.914 | F (9, 36) = 0.3632 | P=0.9451 |
| Residual (random) | 487 | 36 | 13.53 | | |
| Total | 1547 | 49 | | | |

| **Tukey's multiple comparisons test** | **Mean Diff.** | **95.00% CI of diff.** | **Summary** | **Adjusted P Value** |
|---|---|---|---|---|
| D vs. J | -1.35 | -8.948 to 6.249 | ns | 0.972 |
| D vs. I | 1.986 | -5.891 to 9.864 | ns | 0.9086 |
| D vs. PC | 7.804 | 1.700 to 13.91 | * | 0.013 |
| D vs. DC | 10.23 | 5.260 to 15.19 | *** | 0.0005 |
| J vs. I | 3.336 | -0.5851 to 7.257 | ns | 0.104 |
| J vs. PC | 9.154 | 4.411 to 13.90 | *** | 0.0008 |
| J vs. DC | 11.58 | 6.580 to 16.57 | *** | 0.0002 |
| I vs. PC | 5.818 | 1.546 to 10.09 | ** | 0.0088 |
| I vs. DC | 8.239 | 3.117 to 13.36 | ** | 0.0029 |
| PC vs. DC | 2.421 | -1.692 to 6.534 | ns | 0.3463 |

**Fig 3 VipCre-tdT**

**Repeated measures ANOVA summary**

| | |
|---|---|
| Assume sphericity? | No |
| F | 41.74 |
| P value | <0.0001 |
| P value summary | **** |
| Statistically significant (P < 0.05)? | Yes |

| **ANOVA table** | **SS** | **DF** | **MS** | **F (DFn, DFd)** | **P value** |
|---|---|---|---|---|---|
| Treatment (between columns) | 1637 | 4 | 409.3 | F (2.499, 19.99) = 41.74 | P<0.0001 |
| Individual (between rows) | 22.7 | 8 | 2.838 | F (8, 32) = 0.2894 | P=0.9645 |
| Residual (random) | 313.8 | 32 | 9.808 | | |
| Total | 1974 | 44 | | | |

| **Tukey's multiple comparisons test** | **Mean Diff.** | **95.00% CI of diff.** | **Summary** | **Adjusted P Value** |
|---|---|---|---|---|
| D vs. J | -3.945 | -7.792 to -0.09721 | * | 0.0444 |
| D vs. I | -0.4929 | -7.135 to 6.149 | ns | 0.9988 |
| D vs. PC | 10.05 | 3.750 to 16.36 | ** | 0.0037 |
| D vs. DC | 10.95 | 5.781 to 16.13 | *** | 0.0006 |
| J vs. I | 3.452 | -2.950 to 9.853 | ns | 0.4047 |
| J vs. PC | 14 | 9.097 to 18.90 | **** | <0.0001 |
| J vs. DC | 14.9 | 11.84 to 17.96 | **** | <0.0001 |
| I vs. PC | 10.55 | 5.496 to 15.60 | *** | 0.0006 |
| I vs. DC | 11.45 | 6.918 to 15.97 | *** | 0.0002 |
| PC vs. DC | 0.9008 | -2.890 to 4.691 | ns | 0.9168 |

**Fig 3 Somatostatin**

**Repeated measures ANOVA summary**

| | |
|---|---|
| Assume sphericity? | No |
| F | 13.93 |
| P value | 0.033 |
| P value summary | * |
| Statistically significant (P < 0.05)? | Yes |

| ANOVA table | SS | DF | MS | F (DFn, DFd) | P value |
|---|---|---|---|---|---|
| Treatment (between columns) | 282.2 | 4 | 70.56 | F (1.009, 3.027) = 13.93 | P=0.0330 |
| Individual (between rows) | 40.02 | 3 | 13.34 | F (3, 12) = 2.633 | P=0.0978 |
| Residual (random) | 60.8 | 12 | 5.067 | | |
| Total | 383.1 | 19 | | | |

| Tukey's multiple comparisons test | Mean Diff. | 95.00% CI of diff. | Summary | Adjusted P Value |
|---|---|---|---|---|
| D vs. J | 0.2995 | -0.7585 to 1.358 | ns | 0.6219 |
| D vs. I | -1.638 | -4.273 to 0.9967 | ns | 0.1637 |
| D vs. PC | -9.838 | -16.97 to -2.701 | * | 0.0206 |
| D vs. DC | -4.465 | -18.97 to 10.04 | ns | 0.5661 |
| J vs. I | -1.937 | -3.547 to -0.3284 | * | 0.0301 |
| J vs. PC | -10.14 | -16.24 to -4.038 | * | 0.0121 |
| J vs. DC | -4.764 | -18.23 to 8.700 | ns | 0.4716 |
| I vs. PC | -8.2 | -12.96 to -3.439 | * | 0.0109 |
| I vs. DC | -2.827 | -14.88 to 9.224 | ns | 0.737 |
| PC vs. DC | 5.373 | -2.010 to 12.76 | ns | 0.113 |

**Ext Fig 1b duo density**

**Ordinary one-way ANOVA summary**

| | |
|---|---|
| F | 26.16 |
| P value | <0.0001 |
| P value summary | **** |
| Significant diff. among means (P < 0.05)? | Yes |
| R squared | 0.8396 |

| **ANOVA table** | **SS** | **DF** | **MS** | **F (DFn, DFd)** | **P value** |
|---|---|---|---|---|---|
| Treatment (between columns) | 6183080 | 5 | 1236616 | F (5, 25) = 26.16 | P<0.0001 |
| Residual (within columns) | 1181584 | 25 | 47263 | | |
| Total | 7364664 | 30 | | | |

| **Dunnett's multiple comparisons test** | **Mean Diff.** | **95.00% CI of diff.** | **Summary** | **Adjusted P Value** |
|---|---|---|---|---|
| P21 vs. E14.5 | -1297 | -1654 to -939.7 | **** | <0.0001 |
| P21 vs. E16.5 | -1121 | -1461 to -780.6 | **** | <0.0001 |
| P21 vs. E18.5 | -856.5 | -1302 to -411.3 | *** | 0.0001 |
| P21 vs. P0 | -1132 | -1637 to -627.2 | **** | <0.0001 |
| P21 vs. P10 | -309.8 | -754.9 to 135.4 | ns | 0.2459 |

**Ext Fig 1c ileum density**

**Ordinary one-way ANOVA summary**

| | |
|---|---|
| F | 16.57 |
| P value | <0.0001 |
| P value summary | **** |
| Significant diff. among means (P < 0.05)? | Yes |
| R squared | 0.7341 |

| ANOVA table | SS | DF | MS | F (DFn, DFd) | P value |
|---|---|---|---|---|---|
| Treatment (between columns) | 10645289 | 5 | 2129058 | F (5, 30) = 16.57 | P<0.0001 |
| Residual (within columns) | 3855587 | 30 | 128520 | | |
| Total | 14500875 | 35 | | | |

| Dunnett's multiple comparisons test | Mean Diff. | 95.00% CI of diff. | Summary | Adjusted P Value |
|---|---|---|---|---|
| P21 vs. E14.5 | -882.9 | -1490 to -275.4 | ** | 0.0027 |
| P21 vs. E16.5 | -1663 | -2220 to -1106 | **** | <0.0001 |
| P21 vs. E18.5 | -1013 | -1829 to -198.4 | * | 0.0111 |
| P21 vs. P0 | -1373 | -1949 to -796.5 | **** | <0.0001 |
| P21 vs. P10 | -554 | -1162 to 53.51 | ns | 0.0822 |

**Ext Fig 1d PC density**

**Ordinary one-way ANOVA summary**

| | |
|---|---|
| F | 7.077 |
| P value | 0.0004 |
| P value summary | *** |
| Significant diff. among means (P < 0.05)? | Yes |
| R squared | 0.6166 |

| ANOVA table | SS | DF | MS | F (DFn, DFd) | P value |
|---|---|---|---|---|---|
| Treatment (between columns) | 9624083 | 5 | 1924817 | F (5, 22) = 7.077 | P=0.0004 |
| Residual (within columns) | 5983409 | 22 | 271973 | | |
| Total | 15607492 | 27 | | | |

| Dunnett's multiple comparisons test | Mean Diff. | 95.00% CI of diff. | Summary | Adjusted P Value |
|---|---|---|---|---|
| P21 vs. E14.5 | 341.8 | -793.5 to 1477 | ns | 0.8553 |
| P21 vs. E16.5 | -1338 | -2298 to -378.7 | ** | 0.0048 |
| P21 vs. E18.5 | -809.2 | -1871 to 252.7 | ns | 0.1712 |
| P21 vs. P0 | -1225 | -2209 to -242.3 | * | 0.0119 |
| P21 vs. P10 | -412.9 | -1428 to 602.5 | ns | 0.674 |

**Ext Fig 1e duod interstripe**

| **Ordinary one-way ANOVA summary** | |
|---|---|
| F | 3.933 |
| P value | 0.0724 |
| P value summary | ns |
| Significant diff. among means (P < 0.05)? | No |
| R squared | 0.6629 |

| **ANOVA table** | **SS** | **DF** | **MS** | **F (DFn, DFd)** | **P value** |
|---|---|---|---|---|---|
| Treatment (between columns) | 22978 | 3 | 7659 | F (3, 6) = 3.933 | P=0.0724 |
| Residual (within columns) | 11684 | 6 | 1947 | | |
| Total | 34662 | 9 | | | |

| **Dunnett's multiple comparisons test** | **Mean Diff.** | **95.00% CI of diff.** | **Summary** | **Adjusted P Value** |
|---|---|---|---|---|
| P21 vs. E18.5 | 78.48 | -45.22 to 202.2 | ns | 0.21 |
| P21 vs. P0 | 136.6 | 1.105 to 272.1 | * | 0.0485 |
| P21 vs. P10 | 28.28 | -95.42 to 152.0 | ns | 0.8142 |

**Ext Fig 1f ileum interstripe**

| **Ordinary one-way ANOVA summary** | |
|---|---|
| F | 15.79 |
| P value | 0.0002 |
| P value summary | *** |
| Significant diff. among means (P < 0.05)? | Yes |
| R squared | 0.678 |

| **ANOVA table** | **SS** | **DF** | **MS** | **F (DFn, DFd)** | **P value** |
|---|---|---|---|---|---|
| Treatment (between columns) | 42586 | 2 | 21293 | F (2, 15) = 15.79 | P=0.0002 |
| Residual (within columns) | 20222 | 15 | 1348 | | |
| Total | 62807 | 17 | | | |

| **Dunnett's multiple comparisons test** | **Mean Diff.** | **95.00% CI of diff.** | **Summary** | **Adjusted P Value** |
|---|---|---|---|---|
| P21 vs. P0 | 119.7 | 65.41 to 174.0 | *** | 0.0002 |
| P21 vs. P10 | 46.69 | -10.53 to 103.9 | ns | 0.1129 |

**Ext Fig 1f PC interstripe**

**Ordinary one-way ANOVA summary**

| | |
|---|---|
| F | 6.765 |
| P value | 0.0161 |
| P value summary | * |
| Significant diff. among means (P < 0.05)? | Yes |
| R squared | 0.6005 |

| ANOVA table | SS | DF | MS | F (DFn, DFd) | P value |
|---|---|---|---|---|---|
| Treatment (between columns) | 15487 | 2 | 7744 | F (2, 9) = 6.765 | P=0.0161 |
| Residual (within columns) | 10302 | 9 | 1145 | | |
| Total | 25789 | 11 | | | |

| Dunnett's multiple comparisons test | Mean Diff. | 95.00% CI of diff. | Summary | Adjusted P Value |
|---|---|---|---|---|
| P21 vs. P0 | 74.34 | 7.339 to 141.3 | * | 0.0318 |
| P21 vs. P10 | -2.923 | -66.99 to 61.14 | ns | 0.989 |
